# Transient astrocytic Gq signaling underlies remote memory enhancement

**DOI:** 10.1101/753095

**Authors:** Youichi Iwai, Katsuya Ozawa, Kazuko Yahagi, Mika Tanaka, Shigeyoshi Itohara, Hajime Hirase

## Abstract

Astrocytes elicit transient Ca^2+^ elevations induced by G protein-coupled receptors (GPCRs), yet their role *in vivo* remains unknown. To address this, transgenic mice with astrocytic expression of the optogenetic Gq-type GPCR, Optoα1AR, were established, in which transient Ca^2+^ elevations similar to those in wild type mice were induced by brief blue light illumination. Activation of cortical astrocytes resulted in an adenosine A1 receptor-dependent inhibition of neuronal activity. Moreover, sensory stimulation with astrocytic activation induced long-term depression of sensory evoked response. At the behavioral level, repeated astrocytic activation in the anterior cortex gradually affected novel open field exploratory behavior, and remote memory was enhanced in a novel object recognition task. These effects were blocked by A1 receptor antagonism. On the other hand, compelling evidence for astrocytic Ca^2+^-induced diameter changes of arteries was not observed. Together, we demonstrate that GPCR-triggered Ca^2+^ elevation in cortical astrocytes has causal impacts on neuronal activity and behavior.

## Introduction

Interfacing both synapses and blood vessels, astrocytes’ prime functions in the brain have been recognized as the maintenance of extracellular environment and the transfer of energy substrates. Additionally, studies in the recent decade have presented compelling evidence that astrocytes modulate neuronal activity by various mechanisms. Central to the astrocyte-mediated modulation of neuronal activity is Ca^2+^ elevation of astrocytes. While electrically passive, astrocytes elicit large-amplitude cytosolic Ca^2+^ elevations that are triggered by G protein-coupled receptors (GPCRs), particularly the Gq-type, which activate the inositol trisphosphate (IP3) pathway. Amongst Gq-GPCRs, the alpha-1 adrenergic receptor (α1AR) has been identified to be the prevalent receptor for brain-wide astrocytic Ca^2+^ elevations, responding to locus coeruleus (LC) activation in awake mice ^1,2^.

On the vascular side, astrocytic Ca^2+^ elevation has been implicated in the modulation of local cerebral blood flow ^3–5^. However, astrocytic modulation of local cerebral blood flow has been questioned since IP3 receptor type-2 knockout mice (IP3R2-KO), in which large astrocytic Ca^2+^ elevations are diminished, display a similar extent of functional hyperemia ^6,7^ The controversy remains unresolved to date. For instance, recent studies that make use of genetic Ca^2+^ indicators are not in support of astrocytic modulation of blood flow ^8,9^ while another study suggests modulation via pericytes ^10^.

A considerable amount of literature suggests modulation of synaptic transmission and plasticity by astrocytic Ca^2+^ elevation ^11–20^. On the other hand, there are a few studies that report negative results ^21,22^. Moreover, a study that performed a behavioral test battery reported no obvious phenotype in astrocyte-specific IP3R2-KO mice ^23^. Recently, however, the use of synthetic GPCRs (i.e. DREADDs) permitted astrocyte-specific pharmacogenetic activation of Gq signaling. While an initial study that targeted brain-wide astrocytes did not find a phenotype in motor learning ^24^, recent studies reported phenotypes in aversive learning by amygdalar or hippocampal astrocytic activation ^25,26^. However, pharmacogenetic activation of astrocytes inevitably leads to hour-long activation of astrocytes, hence the role of physiological activation of astrocytes in behaving mice has remained unaddressed. Here, we generated transgenic mouse lines in which astrocytes express the optogenetically activated Gq-GPCR Optoα1AR ^27^ that permits transient elevation of astrocytic Ca^2+^ by blue light in inflammation-free conditions.

## Results

### Transgenic mice with selective astrocytic expression of Optoα1AR

We have generated transgenic mice in which Optoα1AR ^27^ is expressed under the control of a BAC-GLT1 promoter ^28^ (**Fig. 1a**). Among the 11 GLT1-Optoα1AR-EYFP founders, a few lines showed astrocyte-specific expression. We examined two lines, #941 and #877, in this study. Line #941 showed ubiquitous expression of Optoα1AR-EYFP in astrocytes throughout the brain (**Fig. 1b; Sup. 1a**), whereas line #877 had expression in a sparse population of astrocytes (**Fig. 1c; Sup. 1b**). EYFP fluorescence in the cortex of line #941 was twice as strong as that of line #877 (195.3 ± 9.0% of 9 strong TG mice relative to 100 ± 7.8% of 10 patchy TG mice; p<0.0001, unpaired t-test). Accordingly, #941 and #877 are referred to as “strong” and “patchy” TG mice, respectively. Immunohistochemistry with astrocytic (S100β), neuronal (NeuN) and microglial (Iba1) markers showed that Optoα1AR-EYFP was expressed in astrocytes (strong TG: 76.9 ± 3.3%, S100β-positive 448 cells, 4 mice; patchy TG: 49.1 ± 2.8%, 476 cells, 4 mice). EYFP expression was negligible in neurons (strong TG: 8.3 ± 1.7%, NeuN-positive 2463 cells, 4 mice; patchy TG: 1.4 ± 0.6%, 2177 cells, 4 mice) and microglia (strong TG: 1.4 ± 0.8%, IbaI-positive 142 cells, 4 mice; patchy TG: 1.6 ± 0.9%, 151 cells, 4 mice) in the cortex (**Sup. 1a and b**). Cortical GFAP and IbaI expressions were generally low in wild type (WT), strong and patchy TG mice (**Sup. 1c,** GFAP, strong TG: 89.5 ± 3.9%, patchy TG: 87.5 ± 2.4%, relative to WT: 100 ± 6.2%. **Sup. 1d,** IbaI, strong TG: 89.2 ± 4.5%, patchy TG: 87.1 ± 3.6%, relative to WT: 100 ± 5.7%, n = 4 mice for all groups). Fine ramified microglial processes were prevalently observed in IbaI immunohistochemistry (**Sup. 1a and b**), suggesting that the expression of the foreign protein does not cause glial inflammation in these TG lines.

**Figure 1.**
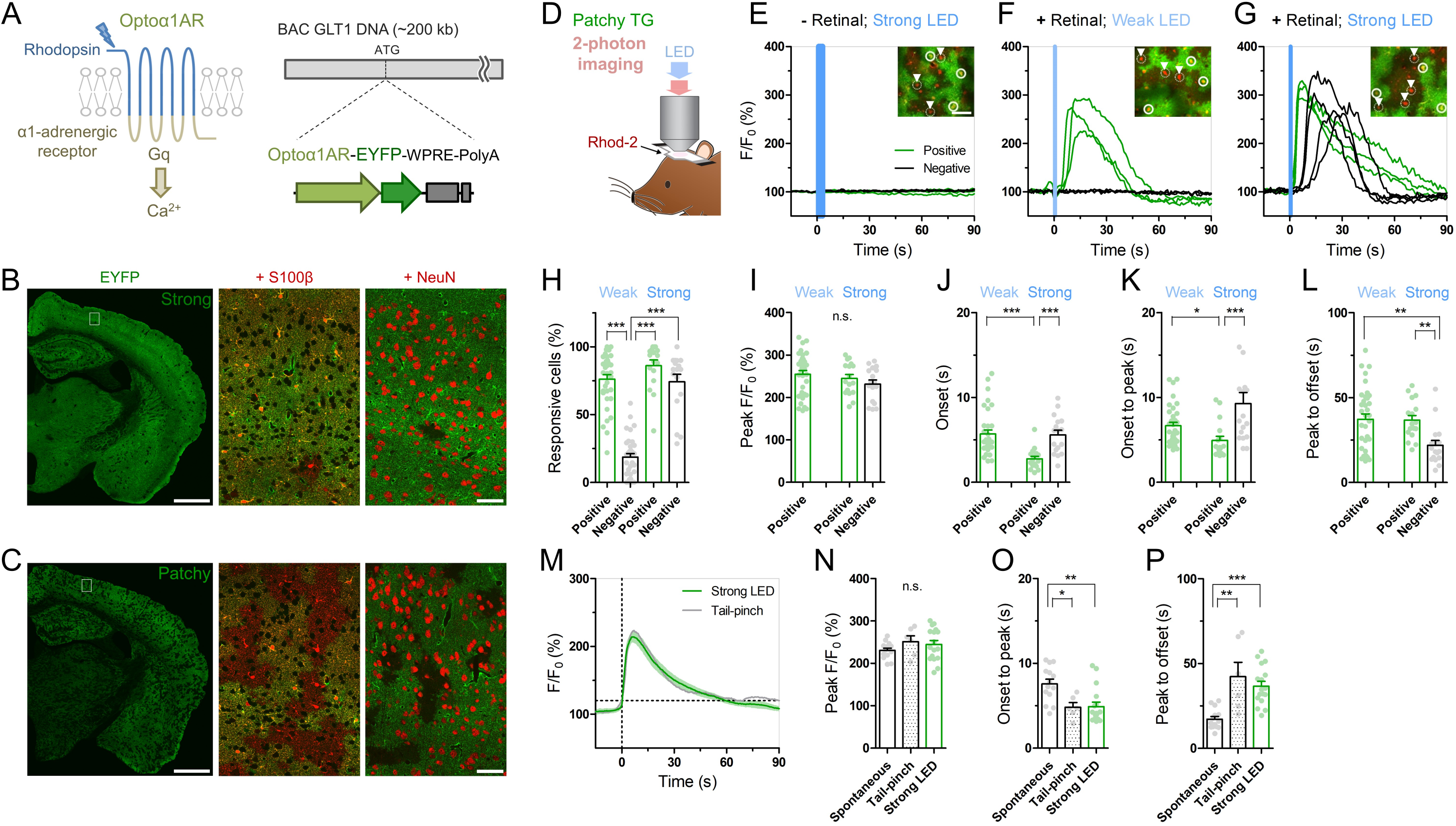
Cis-Retinal supplement is required for reliable Optoα1AR activation by brief illumination *in vivo*. (A) Optoα1AR, optically-activatable Gq-GPCR, is a chimeric molecule of mammalian rhodopsin and Gq-coupled α1 adrenergic receptor ^27^, which induces intracellular Ca^2+^ elevation upon activation. For astrocyte-selective expression, a TG vector was constructed with the BAC GLT1 DNA ^28^. (B) Line #941 (“Strong”) shows intense EYFP fluorescence (green) throughout the brain. High magnification view in white rectangle shows that EYFP is expressed in S100β–positive astrocytes (red). Relatively high EYFP signals are visible in astrocytic somata and endfeet. By contrast, hardly any NeuN signals (red) from neurons overlap with EYFP signals. Scale bar: 1 mm (left), 50 μm (middle and right). (C) Line #877 (“Patchy”) shows visible EYFP fluorescence (green) in a patchy pattern throughout the cortex, hippocampus and striatum. High magnification view in white rectangle shows that EYFP signals colocalize with S100β signals (red) in roughly half of the astrocytes. There are EYFP-negative but S100β-positive domains. Relatively high EYFP signals are visible in astrocytic somata and endfeet. NeuN signals (red) do not overlap with EYFP signals. Scale bar: 1 mm (left), 50 μm (middle and right). (D) Sketch of *in vivo* astrocytic Ca^2+^ imaging with LED illumination in cortical superficial layers of urethane-anesthetized patchy-TG mice. Astrocytes are loaded with the red Ca^2+^ indicator Rhod-2. (E-G) Example plots of *in vivo* astrocytic Ca^2+^ imaging with optical stimulation. Optoα1AR-positive and negative astrocytes (white circles and arrowheads, respectively) are analyzed based on the EYFP expression. Green and black traces correspond to EYFP-positive and -negative astrocytes, respectively. Insets: images of cells analyzed in the respective plots. Scale bars: 50 μm. (E) Astrocytic Ca^2+^ imaging without retinal addition. Strong blue LED illumination (1 mW, 5 s) did not induce Ca^2+^ elevations in all the encircled 6 astrocytes. (F) Astrocytic Ca^2^ imaging with retinal. Weak LED illumination (0.1 mW, 1 s) induced a transient Ca^2+^ increase in EYFP-positive astrocytes, but not in EYFP-negative astrocytes. (G) Astrocytic Ca^2^ imaging with retinal. Strong LED illumination (1 mW, 1 s) induced a rapid Ca^2+^ increase in EYFP-positive astrocytes. Delayed Ca^2+^ elevation was observed in EYFP-negative astrocytes. (H-L) Analysis of Ca^2+^ response in EYFP-positive and -negative astrocytes upon weak or strong LED illumination with retinal addition. Each symbol represents an individual imaging session (Weak LED: 35 sessions, 9 patchy TG mice; Strong LED: 17 sessions, 8 patchy TG mice). (H) Proportion of responsive cells. The weak-negative group was the least responsive (***p<0.001, Tukey’s test after one-way ANOVA), hence the data of this group were not included in the following figures (I to L). (I) Peak amplitude was similar among the remaining 3 groups (p>0.22, one-way ANOVA). (J and K) Onset time and onset-to-peak time were shorter in the strong-positive group (***p<0.001, *p<0.05, Dunn’s test after Kruskal-Wallis one-way ANOVA). (L) Peak-to-offset time was shorter in the strong-negative group (**p<0.01, Dunn’s test after Kruskal-Wallis one-way ANOVA). (M-P) Comparison of spontaneous, tail-pinch-induced, and optogenetically induced (strong illumination) Ca^2+^ response. (M) Mean and SEM trace of optogenetically induced (green) and tail-pinch-induced (gray) Ca^2+^ increase. Time 0 corresponds to onset time, when F/F_0_ reaches 120 %. (N) Peak amplitude was similar among the 3 groups (p> 0.32, one-way ANOVA). (O and P) Onset-to-peak and peak-to-offset times of Ca^2+^ events were similar between the tail pinch and optogenetically induced groups (p>0.05, Tukey’s test after one-way ANOVA) and distinct from spontaneously observed Ca^2+^ events (***p<0.001, **p<0.01, *p<0.05, Tukey’s test after one-way ANOVA). Each symbol represents an individual imaging session (Spontaneous: 14 sessions, 10 TG mice; Tail-pinch: 6 sessions, 4 TG mice).

Next, we performed *in vivo* Ca^2+^ imaging from superficial layers of the cortex in urethane-anesthetized patchy TG mice. Cortical astrocytes were loaded with the red Ca^2+^ indicator Rhod-2 and observed by two-photon microscopy (**Fig. 1d)**. Optoα1AR-positive astrocytes were distinguishable by their EYFP fluorescence, allowing simultaneous investigation of Optoα1AR-positive and -negative astrocytes (**Fig. 1e-g, insets**). Activation of the vertebrate rhodopsin-based GPCRs including the Optoα1AR requires cis-retinal, which is converted to trans-retinal and released upon activation ^29,30^. Consistent with low levels of endogenous cis-retinal in the cortex, blue light illumination did not induce Ca^2+^ elevations in Optoα1AR-positive astrocytes even when power and duration of the blue light were increased (**Fig. 1e**). Next, we repeated the experiment with a supplement of 9-cis-retinal by i.p. injection. Notably, systemic 9-cis-retinal administration enabled reliable photoactivation of Optoα1AR (**Fig. 1f and g; Sup. Movie 1 and 2**). Upon brief blue light illumination (1 s, 0.1 mW) at the surface of the cortex through the objective lens, Optoα1AR-positive astrocytes elevated their Ca^2+^ levels with a delay of 5.71 ± 0.45 s from the onset of illumination (1717 out of 2162 cells, 35 sessions, 9 mice; **Fig. 1f, h and j**). On the other hand, a significantly lower proportion of simultaneously imaged Optoα1AR-negative astrocytes responded to the photostimulation (270 out of 1484 cells, 35 sessions, 9 mice; **Fig. 1f and h**).

Remarkably, more intense illumination (1 s, 1 mW) gave rise to Ca^2+^ elevations in Optoα1AR-negative astrocytes (**Fig. 1g and Sup. Movie 2**). Comparison of Optoα1AR-positive and - negative astrocytes revealed that Optoα1AR-positive astrocytes had a faster onset (2.76 ± 0.30 s vs 5.59 ± 0.54 s; 646 out of 764 cells vs 274 out of 358 cells, 17 sessions, 8 mice; **Fig. 1j**) and rise time (onset to peak: 4.92 ± 0.51 s vs 9.28 ± 1.29 s; **Fig. 1k**). The decay time was also longer in positive astrocytes (peak to offset: 36.7 ± 2.76 s vs 21.8 ± 2.90 s; **Fig. 1l**).

We compared optically evoked astrocytic Ca^2+^ elevations with spontaneous Ca^2+^ elevations which occur with low frequencies under urethane anesthesia ^31,32^. We found that although both magnitudes are similar, optically induced Ca^2+^ elevations are faster in the rise time and slower in the decay time than spontaneous Ca^2+^ events (optically evoked vs spontaneous, peak F/F_0_: 244.8 ± 9.33% vs 230.9 ± 5.17%; onset to peak: 4.92 ± 0.51 s vs 7.58 ± 0.56 s; peak to offset: 36.7 ± 2.76 s vs 17.2 ± 1.61 s; **Fig. 1n-p**). On the other hand, optically induced Ca^2+^ elevations are similar to tail-pinch induced Ca^2+^ events (peak F/F_0_: 251.4 ± 13.7%; onset-peak: 4.85 ± 0.536 s; peak-offset: 42.3 ± 8.47 s; **Fig. 1m-p**), suggesting that the optically-evoked Gq signaling mimics the salient stimuli-evoked *in vivo* event (**Fig. 1m**).

We further investigated the optimal light duration for astrocytic Gq activation. To reliably image astrocytic Ca^2+^ level for a long period, RCaMP was expressed in layer 2/3 astrocytes of strong TG mice by AAV9-hGFAP-RCaMP1.07 (**Sup. 2a**; ^33^). RCaMP in the same astrocytes was repeatedly imaged with varying optical stimulation length (1 mW. 1 s, 3 s or 3 min) before and after systemic retinal injection (**Sup. 2a right**). We confirmed that the retinal addition is indispensable for Optoα1AR activation even with 3-min illumination (**Sup. 2b and d**). With retinal, 1-s illumination was sufficient to elevate the Ca^2+^ levels to saturation, since 3-s illumination only marginally increased the responsive cell number with similar peak amplitudes (**Sup. 2e**). Further, 3-min illumination resulted in a single transient Ca^2+^ elevation with a rather smaller peak amplitude (**Sup. 2c and e**). In contrast, shorter light activation (1 s or 3 s) was repeatable with an interval of 1 min, although the response magnitude was diminished by about three folds from the second stimulus onwards (**Sup. 2f**). Inter-stimulus intervals of 3 minutes and 9 minutes restored the original response magnitude by 83.4% and 116.0%, respectively (**Sup. 2g**).

The following experiments therefore were performed with retinal pretreatment by i.p. injection (see methods) unless otherwise noted.

### Astrocytic Gq signaling does not induce artery dilation

To test whether astrocytic Gq signaling mimicking salient sensory stimulation has a causal impact on local cerebral blood flow, we performed two-photon imaging of the vasculature in the somatosensory cortex of urethane-anesthetized strong TG mice **(Fig. 2a**). The vasculature was labeled by i.v. FITC injection, and astrocytes were loaded with Rhod-2. First, we confirmed that brief forelimb stimulation (2 s) induces penetrating artery dilation in the corresponding primary somatosensory cortex. The arterial cross section expanded immediately after stimulus onset and reached to a peak within three seconds. Thereafter, the arterial cross section area gradually decreased to baseline in five seconds. This response was stereotypical and repeatable (**Fig. 2b**). Sensory stimulation was given at an interval of thirty seconds and the degree of artery expansion did not significantly differ between the first three and the last three trials of the nine consecutive trials (p>0.7, paired t-test). Overall, the relative area increase of arteries in the first 3 seconds after stimulus onset was 12.1±0.9% (17 arteries, 5 mice, **Fig. 2c**). On the other hand, arterial area increase in the hindlimb somatosensory cortex was negligible (1.9±0.6%, 4 arteries, 2 mice), suggesting that the arterial dilation was locally induced. Astrocytic endfoot Ca^2+^ elevations were only occasionally observed after sensory stimulation (occurrence probability = 10/100 = 10 %; 10 endfeet, 3 mice). On average, endfoot Ca^2+^ increase (F/F_0_) of 12±5% was observed in the averaged plot with a latency of several seconds (**Fig. 2d**).

**Figure 2.**
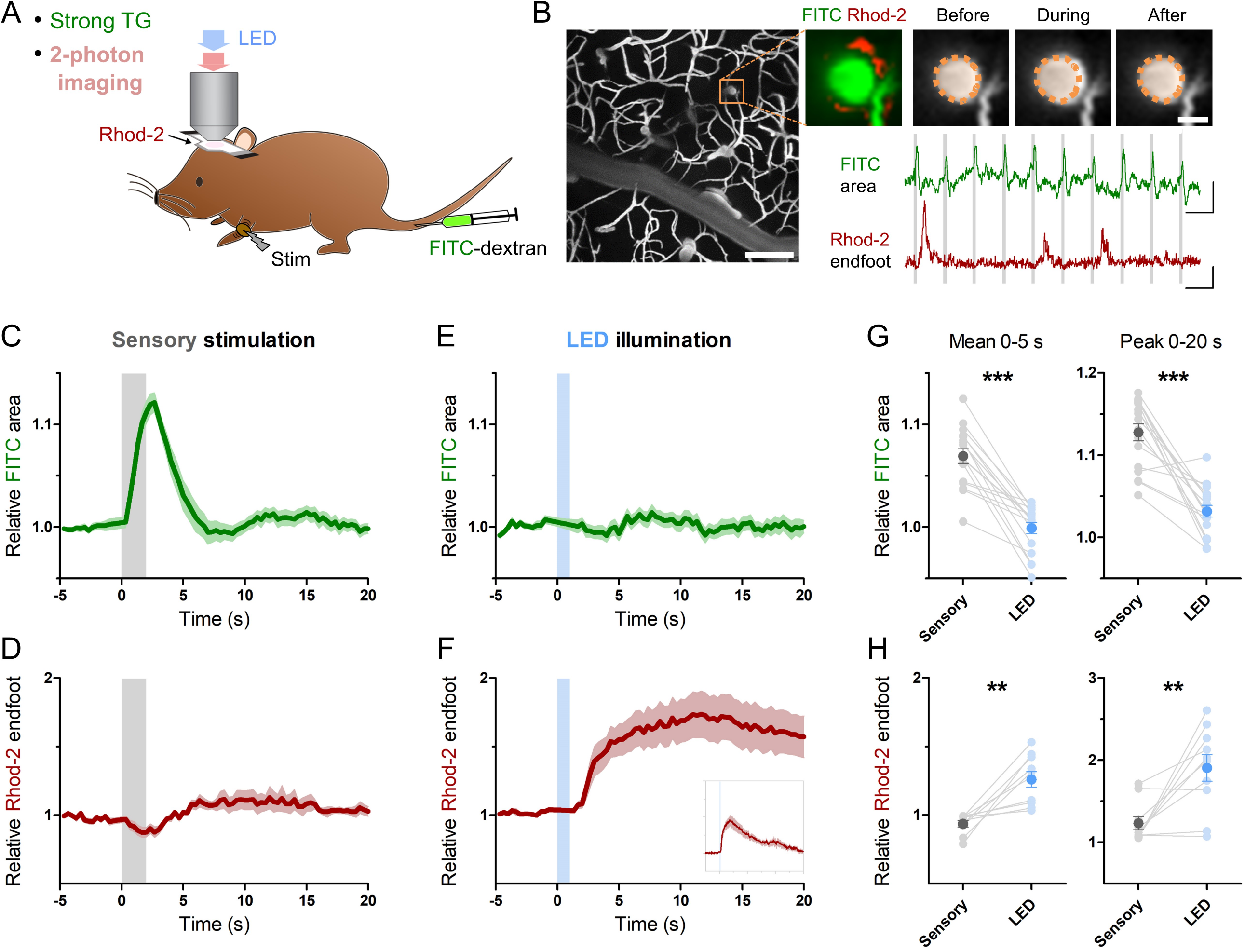
Transient Optoα1AR activation did not induce artery dilation *in vivo*. (A) Two-photon vasculature imaging from layer 2/3 somatosensory cortex of urethane-anesthetized strong TG mice upon LED illumination. Retinal was supplemented prior to imaging. Blood vessels were labeled by i.v. injection of FITC-dextran and astrocytes were loaded with the Rhod-2 Ca^2+^ indicator. LED illumination (1 mW, 1 s) or sensory stimulation (duration 2 s, interval 30 s) were delivered through the objective lens or on the forelimb, respectively. (B) The low-power view shows pattern of FITC-filled blood vessels. The high-power view of the orange rectangle region contains an FITC-filled penetrating arterial cross section (green) and a Rhod-2-loaded astrocytic endfoot (red). The artery expanded during the sensory stimulation. The semi-lucent orange area represents the arterial area before sensory stimulation. The arterial area expanded reliably at every sensory presentation, whereas endfoot Ca^2+^ signal increased only occasionally. Scale bars: 100 μm (low-power image); 10 μm (high-power image); 20 % (FITC area trace), 300 % (Rhod-2 F/F_0_ trace), 30 s (time axis). (C) Arterial cross-section area rapidly increased immediately after the sensory stimulation onset (17 arterial cross sections from 5 strong TG mice). SEM is shown by the shaded region. (D) Averaged endfoot Ca^2+^ signal shows marginal changes after the sensory stimulation (10 endfeet from 3 strong TG mice). (E) Arterial cross-section area of the same data set as in (C) did not increase upon the LED illumination. (F) Endfoot Ca^2+^ signal of the same dataset as in (D) strongly increased after the LED illumination. Insets: Endfoot Ca^2+^ change by LED illumination is shown in full in an expanded time scale. Abscissa: -15 s to 90 s (i.e. the same as in Fig 1M), where time 0 corresponds to the onset of LED illumination. Ordinate: -0.5 to 3.0 F/F_0_. (G) Mean and peak values of arterial cross section area are plotted for the time periods 0–5 s and 0–20 s (time 0 is the onset of sensory stimulation). Both measures are significantly larger than those of LED illumination (p<0.001, p<0.001, paired t-test). (H) Mean and peak values of endfoot Ca^2+^ signal are plotted for the time periods 0–5 s and 0–20 s (time 0 is the onset of LED illumination). Both measures are significantly higher than those of sensory stimulation (p<0.002, p<0.006, paired t-test).

Having demonstrated reliable functional hyperemia in arteries of the somatosensory area, we examined if optically evoked Gq-driven astrocytic Ca^2+^ increase impacts vasomodulation in the same set of arteries. As in **figure 2a**, a brief illumination of blue light (1 s, 1 mW) through the objective lens was given while arterial cross sections and astrocytic Ca^2+^ were imaged. We observed that arterial cross section areas remained unchanged despite optical stimulation (**Fig. 2e**), while Ca^2+^ elevations at astrocytic endfeet were reliably evoked (**Fig. 2f**). The lack of arterial cross section change was in stark contrast with sensory-induced hyperemia (**Fig. 2g**), in spite of significant Ca^2+^ increases in the astrocytic endfeet by optogenetic Gq-GPCR stimulation (**Fig. 2h**). Taken together, these experiments dissociated the role of astrocytic Ca^2+^ elevations from sensory-driven arterial dilation.

### Astrocytic Gq signaling inhibits neuronal activity

To investigate the effect of astrocytic Gq activation on neuronal activity, we monitored neuronal Ca^2+^ activities in layer 2/3 of the somatosensory cortex using AAV1.Syn.NES-jRGECO1a.WPRE.SV40 which allows selective expression of the Ca^2+^ probe in neurons ^34^. Brief LED illumination (1 s, 1 mW) in awake TG mice significantly reduced neuronal Ca^2+^ activity quantified as the standard deviation (std) of somatic F/F_0_ jRGECO1a (**Fig. 3a** and **Sup. Movie 3**). This somatic activity reduction was detectable during the first 1 min (**Fig. 3c**; jRGECO1a F/F_0_ std relative to pre-LED 1 min period: 83.0±4.3%; p<0.008) and continued for an additional minute (**Fig. 3c**; 88.2±3.3%; p<0.02). Similar Ca^2+^ activity reduction was also detected in the neuropil (**Fig. 3a** and **c**; post-LED 1 min period: 84.1±5.1%; p<0.03; post-LED 1-2 min period: 90.9±3.2%; p<0.03). These suppressive effects could not be attributed to possible photodamage by the two-photon laser or a sensory processing of the LED light, as WT mice did not display such a reduction (**Fig. 3b**; soma, post-LED 1 min period: 98.0±5.0%; p>0.70), which was significantly different from TG (p<0.05, unpaired t-test).

**Figure 3.**
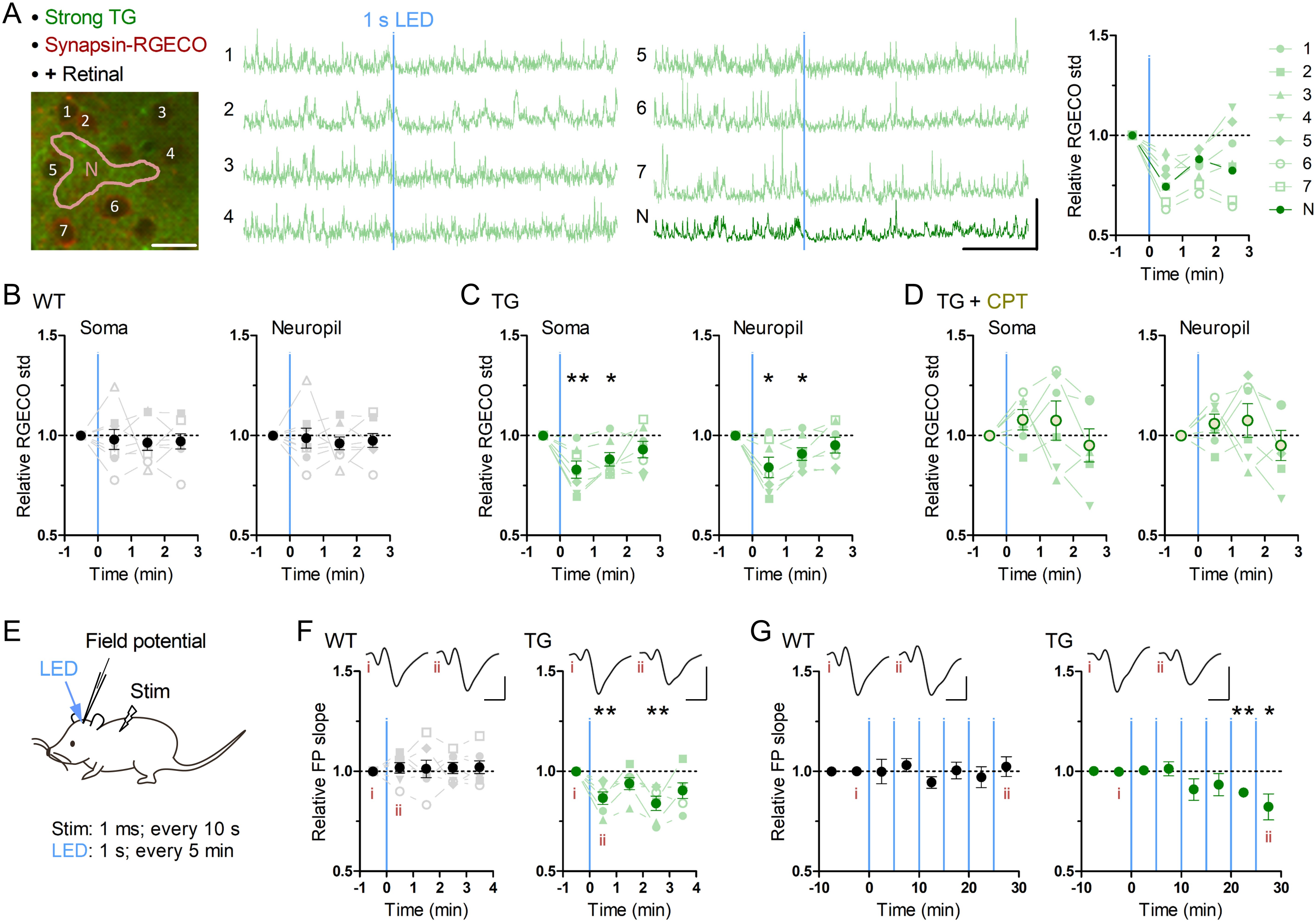
Brief astrocytic Gq activation suppresses neuronal activity. (A-D) Neuronal Ca^2+^ imaging from somatosensory cortex layer 2/3 in awake mice with optogenetic induction of Gq signaling in astrocytes. (A) Representative two-photon image of somatosensory cortex of a strong TG mouse expressing jRGECO1a (RGECO) in neurons by AAV-Syn-jRGECO1a (left). RGECO F/F_0_ from the labeled somata (1-6) and neuropil (N) decreased rapidly after 1 s LED illumination (middle). Ca^2+^ activity, measured as the standard deviation (std) of RGECO F/F_0_, decreased in the first and the second 1 min after LED illumination (right). Scale bars: 20 μm (micrograph); 100 % F/F_0_ and 1 min (traces). (B) Ca^2+^ activity of neuronal somata and neuropil in WT mice did not change after LED illumination (p>0.70 and p>0.80, paired t-test, 1 min after LED illumination vs 1 min before LED illumination, 8 mice). (C) Ca^2+^ activity of neuronal somata and neuropil in TG mice decreased in the first and second minutes after LED illumination (first minute: p<0.008 and p<0.03; second minute: p<0.02 and p<0.03, paired t-test vs 1 min before LED illumination, 7 mice). (D) Adenosine A1R antagonist CPT blocked Optoα1AR-induced neuronal Ca^2+^ activity decrease in somata and neuropil (p>0.18 and p>0.25, paired t-test, 1 min after LED illumination vs 1 min before LED illumination, 6 mice). (E-G) Sensory evoked field potential (FP) recording in somatosensory cortex layer 2/3 of shallowly anesthetized mice upon LED illumination. (E) FP response was evoked by sensory stimulation to the trunk (duration 1 ms, interval 10 s) before and after brief LED illumination (1 mW, duration 1 s). Six optical stimulations (5 min interval) were performed in a session. (F) LED time-triggered averaging of FP slope shows a reduction of sensory evoked response after astrocytic Gq activation in the first 1 min (p<0.008, paired t-test vs 1 min before LED illumination, 6 TG mice). WT mice did not show a significant change in FP slope (p>0.51, paired t-test vs 1 min before LED illumination, 7 WT mice). This reduction in TG mice was detectable 3 min after LED illumination (p<0.008, paired t-test vs 1 min before LED illumination, 6 TG mice). Insets: averaged FP traces from a representative mouse, with the left and right traces averaged within 1 min before and 1 min after LED illumination, respectively. Scale-bars: 200 μV and 20 ms. (G) In the 30 min recording, evoked FP slope gradually decreased in TG mice (20-25 min and 25-30 min periods: p<0.004 and <0.05, paired t-test vs 0-10 min before LED illumination, 6 TG mice), while that in WT mice did not change throughout the 30 min period (p>0.1, paired t-test vs 0-10 min before LED illumination, 7 WT mice). Insets: averaged FP traces from a representative mouse, with the left and right trace averaged within the 5 min period before the first LED illumination and the 5 min period after the last LED illumination, respectively. Scale bars: 200 μV and 20 ms.

Among the molecules released by astrocytic activation, adenosine exerts inhibitory effects through the adenosine A1 receptor (A1R) ^25,35,36^. When the A1R antagonist, CPT, was applied, this astrocytic Optoα1AR-induced neuronal suppression disappeared and a trend for neuronal activation was observed (**Fig. 3d**; soma post-LED 1 min period: 107.9±5.1%; p>0.18), which was significantly different from that of TG mice without CPT (p<0.004, unpaired t-test). These results suggest that transient astrocytic Gq activation rapidly suppresses spontaneous neuronal activity in awake condition via A1R. Of note, similar experiments in urethane-anesthetized strong TG mice resulted in a milder decrease of neuronal activity (**Sup. 3a**) presumably due to lower basal Ca^2+^ activity (**Sup. 3b**).

To examine the effect of astrocytic Gq activation on sensory evoked neuronal activity, we performed *in vivo* field potential (FP) recording from somatosensory cortex layer 2/3 under shallow isoflurane anesthesia (~0.8%) (**Fig. 3e**). After stable FP response for sensory stimulation on the trunk was obtained, brief LED illuminations were delivered from the pial surface above the recording site through optical fiber (φ = 0.2 mm, 1 mW, duration 1 s, interval 5 min, 6 times). As a result, evoked FP slope was decreased after the LED illumination in TG mice, while it was unchanged in WT mice (**Fig. 3f**). This reduction was rapidly expressed and lasting (**Fig. 3f**; 86.7±3.1%, 93.9±3.0%, 84.0±3.6% and 90.0±3.9% during the first, second, third and fourth 1 min after LED; p<0.008, >0.09, <0.008 and >0.05, paired t-test). Furthermore, in the course of a 30-min recording, the reduction of evoked FP slope in TG gradually built up (**Fig. 3g**; 89.4±2.0% and 82.3±6.5% during 20-25 min and 25-30 min after the first LED; p<0.004 and <0.05, paired t-test). These results suggest that transient astrocytic Gq activation inhibits evoked synaptic response and repeated astrocytic Gq activations lead to synaptic depression.

### Behavioral impact of astrocytic Gq signaling

Although multiple studies show that noradrenergic Gq signal simultaneously activates astrocytic Ca^2+^ elevation in wide cortical regions ^1,2,37^, its *in vivo* roles remain unknown. To understand how global astrocytic Gq signaling affects mouse behavior, we performed a set of behavioral experiments with Optoα1AR TG mice whereby photostimulation was made using a head-mounted wireless LED device ^38,39^. A blue LED (φ = 3.1 mm) was placed on each side of the thinned skull above the anterior cortex, where dense noradrenergic inputs are projected in the upper layers ^40^.

A single exposure of blue light for 3 s to wide anterior cortical areas did not result in obvious immediate behavioral changes in strong TG mice. For instance, there was no sign of arousal from sleep or falling to sleep by the optical stimulation. We next investigated the exploratory behavior of mice in a novel open field for 45 min while intermittently illuminating the anterior cortical areas (duration 3 s, interval 3 min, 15 times; **Fig. 4a**). As demonstrated in a single-animal example in **figure 4a**, the TG mouse had a sign of lower exploratory behavior than the WT mouse. To check the level of anxiety, time in center domain was quantified, and there was not a significant difference between WT and TG mice in any of the trichotomized time intervals (**Fig. 4b**). Locomotion was consistently lower in optically stimulated TG mice throughout the course of the open field test and the difference from WT mice became more significant in the middle and final 15 min periods (**Fig. 4c and d**). Analysis of immobile time shows that TG mice gradually develop immobility and the difference from WT mice becomes distinct in the last 20 min period (**Fig. 4e**). Time-averaged analysis of locomotion with respect to LED illumination indicates that the transient astrocytic Gq activation reduced the locomotion activity rapidly. This reduced activity continued until the next astrocytic Gq activation (**Fig. 4f**). Therefore, the astrocytic Gq signal-triggered reduction of locomotion accumulated at every LED illumination, resulting in larger locomotion differences detected in later periods (**Fig. 4c and d**). Notably, when the A1R antagonist DPCPX was applied at the dosage that does not affect open field locomotion in WT mice ^41,42^ (1 mg/kg), the reduced locomotion in the strong TG mice by LED illumination was reverted (**Fig. 4d and f**). Patchy TG mice showed a similar trend for reduced locomotion activity, although the degree of reduction was smaller (**Sup. 4**), suggesting that thorough activation of astrocytic Gq signaling is required for the decreased locomotion. Consistently, insufficient Gq activation without retinal supply could not induce the locomotion decrease (**Sup. 5**).

**Figure 4.**
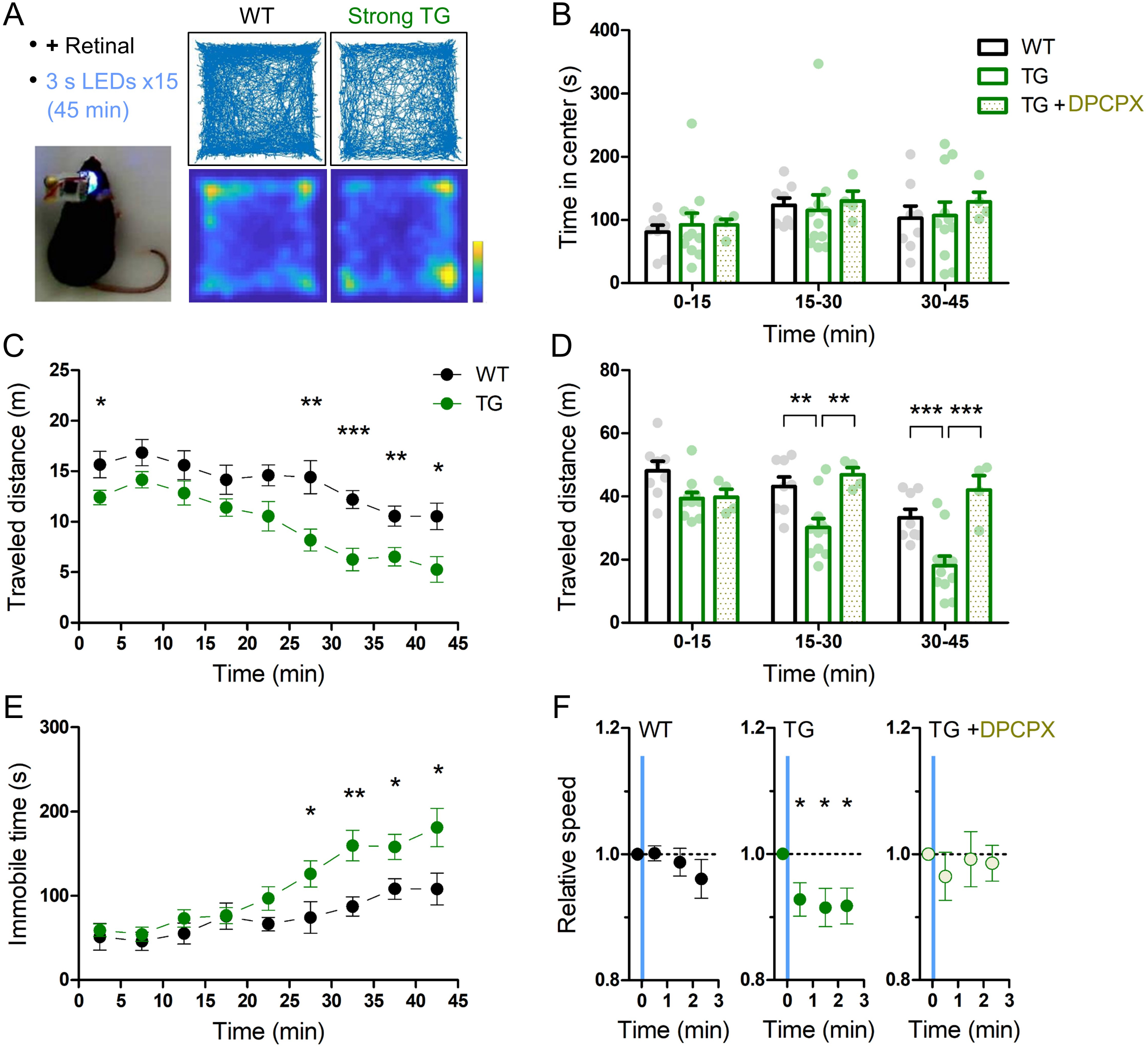
Transient astrocytic Gq activation in the anterior cortex decreases locomotion in a novel open-field. (A) Anterior cortical areas of freely behaving mice were illuminated by a wireless LED device. Representative trajectories and occupancy maps during 45 min open-field behavior of a WT or a strong TG mouse. Both mice received LED illuminations (duration 3 s, interval 3 min, 15 times) with retinal pre-treatment (i.p.). The TG mouse traveled a shorter distance, while spending a similar length of time in the center zone as the WT mouse. Color bar: 15 s. (B) Time in the center zone was not significantly different between experimental conditions and between 15 min periods (p>0.75 and p>0.17, two-way ANOVA, 8 WT mice, 11 strong TG mice vs 4 strong TG mice with DPCPX). (C) TG mice gradually exhibited shorter traveled distances (***p<0.001, **p<0.01, *p<0.05, unpaired t-test, 8 WT mice vs 11 strong TG mice). (D) Traveled distance in TG mice was significantly shorter in 15-30 min and 30-45 min, which was reinstated by DPCPX injection (**p<0.01, ***p<0.001, Bonferroni test after two-way ANOVA). (E) TG mice gradually increased immobile time (*p<0.05, **p<0.01, unpaired t-test, 8 WT mice vs 11 strong TG mice). (F) LED-triggered averaging indicates a rapid and lasting decrease of locomotion in TG mice. Locomotion speed of TG mice was significantly reduced in 0-60 s, 60-120 s and 120-160 s after LED illumination in comparison with that in 0-20 s before LED illumination (p<0.03, p<0.02 and p<0.02, paired t-test). Locomotion speed of WT mice or TG mice with DPCPX did not change significantly after LED illumination (p>0.2 or p>0.4, paired t-test).

Next, we examined the influence of astrocytic Gq signaling on memory. Anterior cortical areas including the prefrontal cortex have been shown to regulate memory acquisition and maintenance ^43–45^, which could be mediated by noradrenergic input ^46^. To evaluate the involvement of astrocytic Gq activation in working memory, we performed the Y-maze test. Both strong TG and WT mice freely explored in the Y-maze for 15 min while receiving blue LED stimulation (duration 3 s, interval 3 min, 5 times; **Fig. 5a**). The probability of the correct arm entry (i.e. the entry to the arm different from the current and immediate prior ones) did not change significantly between TG and WT mice, although its variability was higher in TG mice (**Fig. 5b**). The number of arm entries, total distance traveled and immobile time were also similar between TG and WT mice (**Fig. 5c-e**). These results indicated that the transient astrocytic Gq signal activation in anterior cortical areas does not affect working memory or alter the short-term exploratory behavior in the Y-maze.

**Figure 5.**
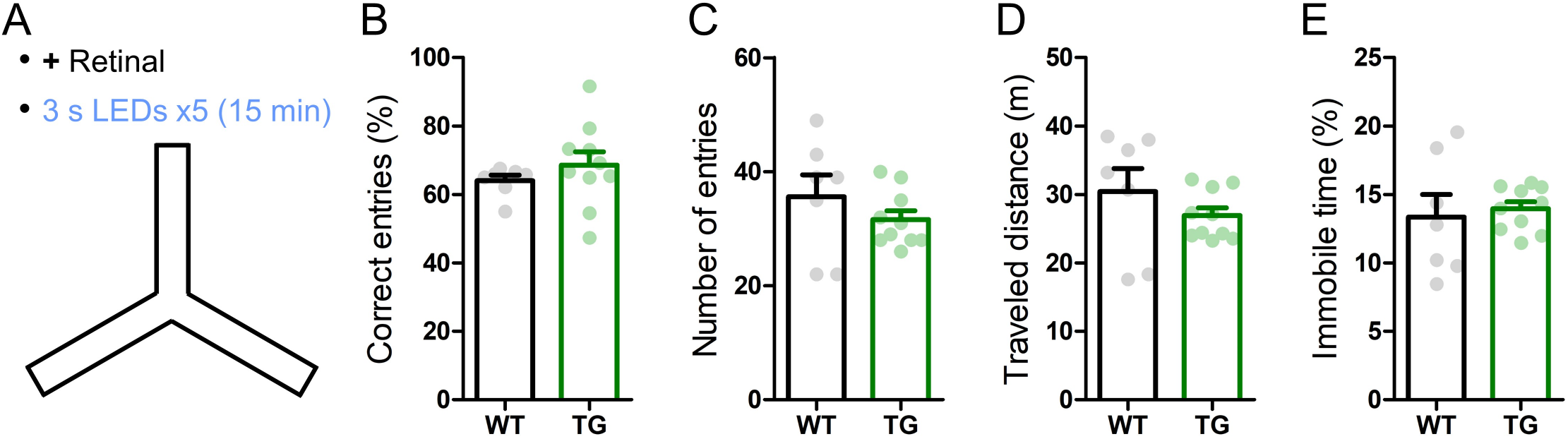
Transient astrocytic Gq activation in the anterior cortex does not affect short-term memory in a Y-maze. (A) WT and strong TG mice were pre-treated with retinal and were put in a Y-maze for 15 min with transient LED illuminations (duration 3 s, interval 3 min, 5 times) delivered. (B) Percentage of correct arm entries (unique triplets) was not significantly different between WT and TG mice (p>0.30, Welch’s t-test, 7 WT mice vs 10 strong TG mice), but the variance was higher in TG mice (p<0.02, F-test). (C, D and E) Number of arm entries, traveled distance and immobile time did not differ between WT and TG mice (p>0.37, p>0.35 and p>0.73, Welch’s t-test; p<0.05, p<0.02 and p<0.02, F-test).

We then tested long-term memory by performing the novel object recognition test. On day 1, anterior cortical areas were illuminated with LED (duration 3 s, interval 3 min, 4 times) during the object familiarization period (training period), whereby two identical objects located apart were exposed for 10 min in a behavior chamber. When one of the pre-exposed objects was replaced with a novel object 1 day later (**Fig. 6a left**), WT and strong TG mice similarly spent a longer time contacting the novel object relative to the familiar object (**Fig. 6a middle**). Notably, when object replacement was done 14 days later (**Fig. 6a left**), TG mice still retained the novel object preference, whereas WT mice lost the preference (**Fig. 6a right**). Even if the duration of each LED illumination was increased from 3 s to 30 s (**Fig. 6b**), TG mice showed a similar novel object preference 14 days later, suggesting that 3 s activation is sufficient to achieve the plateaued enhancement in object memory retention. Of note, LED illumination of TG mice without retinal supply could not significantly induce this 14-day memory retention (**Fig. 6c**), consistent with insufficient Gq activation. Moreover, LED illumination of TG mice with DPCPX treatment did not result in 14-day memory retention, although the novel object recognition after 1 day was normal (**Fig. 6d**). These experiments suggest that transient activation of astrocytic Gq signal does not affect memory acquisition but enhances memory that lasts for more than two weeks through A1R activation.

**Figure 6.**
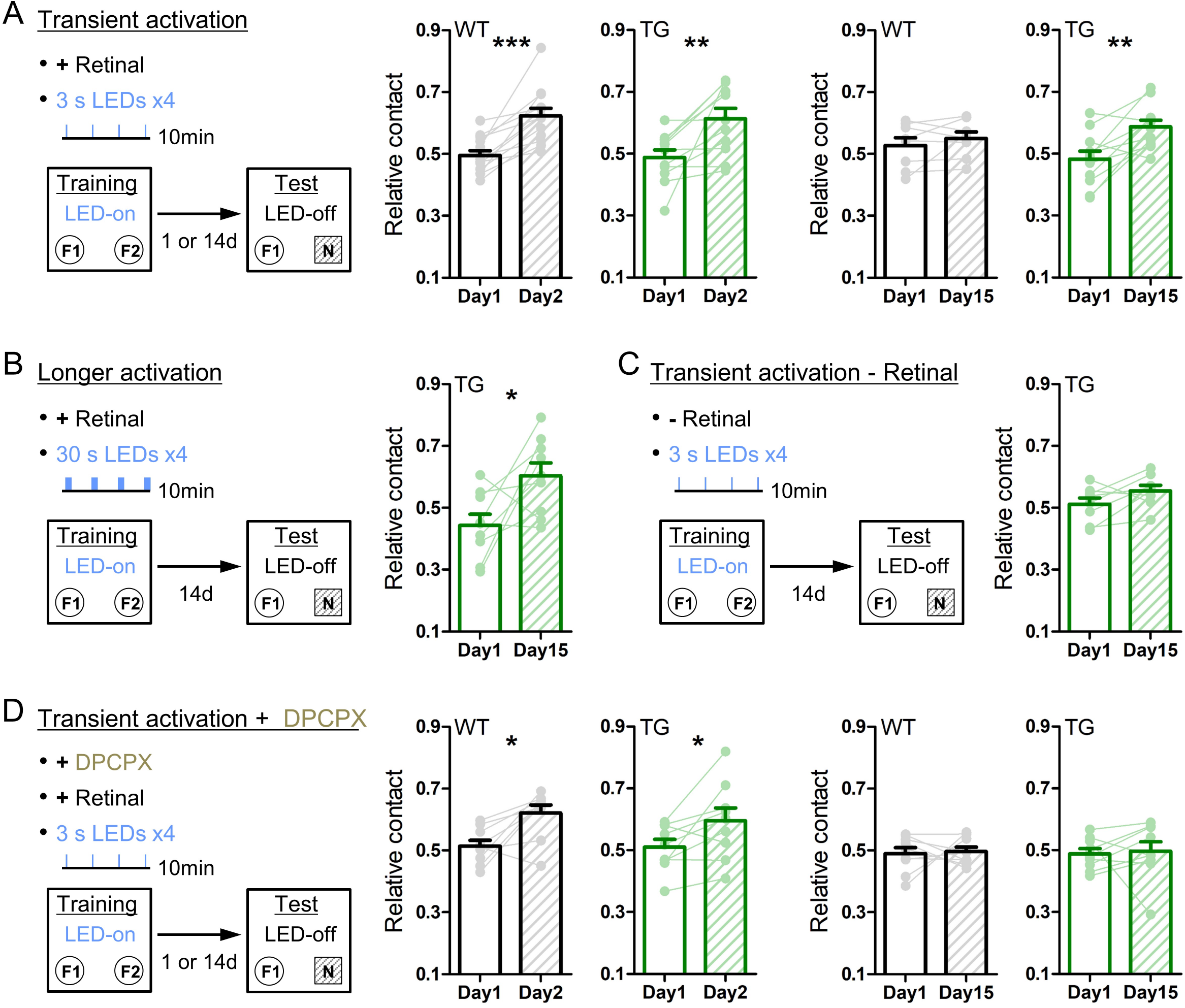
Transient astrocytic Gq activation in the anterior cortex enhances long-term object recognition memory. (A) Transient activation protocol. On the training day, WT or strong TG mice were placed in an open-chamber for 10 min, where two identical objects (F1 and F2) were placed apart. Mice were pre-treated with retinal. Transient LED illuminations (duration 3 s, interval 3 min, 4 times) were delivered during the training period. On the test day, (1 day or 14 days after training), mice were exposed to one of the pre-familiarized objects (F1) and a novel object (N) for 10 min without LED illumination. In the 1-day test, both WT (black) and TG (green) mice similarly increased relative contact time to the novel object. (p<0.0002 and p<0.08 paired t-test vs relative contact time to F2 object in the training, 13 WT mice and 11 TG strong mice). In the 14-day test, TG mice still retained the novel object preference (p<0.005, paired t-test, 11 strong TG mice), whereas WT mice did not (p>0.31, paired t-test, 8 WT mice). (B) Longer activation protocol. The training procedure is the same as in A, except for longer LED illuminations (30 s) delivered. In the 14-day test, TG mice showed the novel object preference (p<0.04, paired t-test, 9 strong TG mice). (C) Transient activation without retinal pre-treatment. The training procedure is the same as in A, except for pre-injection of vehicle instead of retinal. In the 14-day test, TG mice did not show the significant novel object preference (p>0.1, paired t-test, 8 strong TG mice). (D) Transient activation with DPCPX protocol. The training procedure is the same as in A, except for additional adenosine A1R antagonist DPCPX pre-treatment. In the presence of DPCPX, the transient LED illuminations did not induce the novel object preference 14 days later (p>0.82 and p>0.82, paired t-test, 9 WT mice and 9 strong TG mice), whereas novel object preference was expressed 1 day after familiarization (p<0.02 and p<0.04, paired t-test, 9 WT mice and 9 strong TG mice).

Previous studies showed that systemic injection of noradrenaline (NA) immediately after training enhances long-term object recognition memory ^47,48^. To test if astrocytic Gq activation induces similar effects, we photo-stimulated TG mice immediately after training (duration 3 s, interval 3 min, 4 times; **Sup. 6a**) and found that this post activation was also effective for memory enhancement 14 days later (**Sup. 6a right**). Unlike prolonged exposure to an open environment (**Fig. 4**), neither locomotion nor object contact time differed significantly during the 10 min training period across genotypes, illumination durations or drug applications (**Sup. 6b and c**). Furthermore, the memory-enhancing effect is not due to an avoidance behavior from familiarized objects which were associated with astrocytic Gq activation, as the conditioned place preference test suggests that astrocytic Gq activation did not change environmental preference to activated side or non-activated side (**Sup. 7**).

## Discussion

The identification of the optogenetic stimulation conditions for the Optoα1AR TG mice allowed causal assessment of physiological Gq signaling in astrocytes for the first time. While Gq activation in astrocytes did not affect arterial diameter, it transiently inhibited local neuronal activity via adenosine A1 receptor, induced depression of evoked response when paired with sensory stimuli, and reduced locomotor activity. Furthermore, transient astrocytic Gq signaling enhanced long-term memory (LTM) while short-term memory (STM) was not affected.

### Astrocytic Gq signaling is not involved in arterial dilation

We demonstrated that sensory stimulation causes artery dilation and occasional astrocytic endfoot Ca^2+^ elevation. However, optogenetically triggered Ca^2+^ elevations via Optoα1AR in the same set of perivascular astrocytes did not result in vessel dilation (**Fig. 2**). Cortical hyperemia has been demonstrated to be actuated by smooth muscle cells ^49^ and capillary dilation precedes that of upstream arteries ^50^. While we did not examine capillary diameters in this study, the lack of the arterial expansion argues against the involvement of astrocytic Gq signal-induced Ca^2+^ elevation in the control of arterial diameter in sensory-evoked functional hyperemia. Neuronally evoked capillary dilatation may be controlled by endfoot Ca^2+^ entry ^51^ possibly through P2X channels ^10^.

### Astrocytic Gq signaling-induced neuronal inhibition

Effects of astrocytic Gq activation on neuronal activity remains controversial. For instance, in a recent pharmacogenetic study ^26^ and an optogenetic study ^52^, astrocytic Gq activation induced long-term potentiation (LTP) in hippocampal CA1 slices, but an earlier pharmacogenetic study reported no effects ^21^. Whereas these studies did not report *in vivo* effects on astrocytic and neuronal activity, we showed rapid reduction of neuronal activity and depression of evoked response after astrocytic Gq activation (**Fig. 3**), in line with the recent monkey prefrontal cortex study with α1AR agonists ^53^. Aside from the preparation differences, one important aspect of the current study is the use of transgenic mice that minimizes concerns on tissue inflammation caused by virus-injection and slice preparation. Indeed, acute astrogliosis alters multiple plasticity-related molecules ^54^, which potentially impact neuronal activity through reactive astrocytes ^55,56^.

Cholinergic activation elevates astrocytic Ca^2+^ and promotes cortical and hippocampal synaptic potentiation when combined with sensory stimulation in anesthetized rodents ^11,16,57^. Moreover, noradrenergic activation via transcranial direct current stimulation induces synaptic potentiation in the cortex ^20^. The apparent difference of plasticity expression may be explained by the manner whereby GPCR is activated. Volume-transmitted neuromodulators act on neuronal and astrocytic GPCRs, both of which may be required for the synaptic potentiation. Likewise, these neuromodulators also activate Gs and/or Gi/o signaling, that may also play an essential role. Importantly, the current study selectively stimulates astrocytic Gq signaling in a temporally defined manner, thereby dissecting specific roles of the signaling in the neural circuit.

### Astrocytic Gq signaling-induced behavioral changes

We found that transient astrocytic Gq activation in wide anterior cortical areas gradually attenuated locomotion in a novel open field (**Fig. 4**), which is consistent with its neuronal suppressive effects (**Fig. 3**). While noradrenergic activation is generally thought to promote arousal, a study showed that LC noradrenergic neuronal discharge pattern modulates locomotor activity: Phasic high-frequency bursts decrease locomotor activity, and tonic low-frequency discharges increase locomotion ^58^. These modes of LC discharge may activate distinct noradrenergic receptors: tonic discharge might preferentially activate the high-affinity α2AR, whereas phasic discharge may additionally activate the low-affinity α1AR and βARs. A previous pharmacogenetic study of systemic astrocytic Gq activation reported reduced locomotor activity and proposed that sustained Gq signaling in the autonomic nervous system might underlie this effect ^24^. Our study indicates that transient astrocytic Gq activations in the anterior cortical area, which includes motor areas, are sufficient to reduce locomotion presumably by inhibition of the neuronal activity.

### Astrocytic Gq signaling-induced long-term memory enhancement

The absence of effects on STM by the astrocytic Gq signal activation in our study (**Fig. 5**) contrasts with earlier studies that showed an impairment of spatial STM by infusion of α1AR agonists in the prefrontal cortex ^59,60^ or an enhancement of STM in T-maze by pharmacogenetic activation of hippocampal CA1 ^26^. This inconsistency could be due to a combination of multiple factors including activation area (global anterior cortex vs local hippocampal CA1), activation mode (transient vs sustained), and astrocytic Ca^2+^ response magnitude. For the last point, our optogenetic activation results in Ca^2+^ elevations similar to natural astrocytic responses (**Fig. 1**), whereas other studies did not address this issue.

We showed that the astrocytic Gq signal activation did not affect object recognition memory retrieval one day after learning, whereas it enhanced memory two weeks after (**Fig. 6**). In contrast, the pharmacogenetic astrocytic Gq signal activation has been reported to enhance 1-day memory in a contextual fear conditioning task ^26^. Adamsky and colleagues also showed that continuous 5-min illumination of astrocytic Optoα1AR (90% duty cycle) similarly induced a 1-day memory enhancement; however, comparison with our result is difficult due to the lack of description for *in vivo* astrocytic Ca^2+^ elevation and cis-retinal supplement, aside from the learning-paradigm difference. Another recent study showed that 3-min illumination of melanopsin expressed in CA1 astrocytes induced a memory enhancement in an object place test probed 30-min after association ^52^. However, direct comparison to our result is difficult due to the lack of *in vivo* activation data as well as the concern that the melanopsin (Opn4-human) used in the study activates the Gi/o pathway in addition to the Gq pathway ^61^. Our study highlights that transient astrocytic Gq activation promotes remote memory that lasts over weeks and is in line with Lee et al. ^62^ that reported impairment of 2-week object recognition memory in mice deficient of astrocytic vesicular release. Considering previous studies describing astrocytic vesicular release-mediated increases of extracellular adenosine ^35,63^, the enhancement of long-term object memory is potentially mediated by purinergic gliotransmission.

### Astrocytic Gq signaling-mediated novelty detection

LTM enhancement by brief astrocytic activation during or shortly after object familiarization (**Fig. 6, Sup. 6**) is reminiscent of previous studies whereby NA was systemically administered immediately after object familiarization ^47,48^. These results are in line with the “behavioral tagging” hypothesis ^64,65^, which is a behavioral analog of the synaptic tagging and capture theory ^66,67^: ordinary training sets tags on relevant synapses, and a novel experience induces the synthesis of plasticity-related proteins (PRPs) that are captured at the tagged synapses for memory consolidation (**Sup. 8**). Previous studies have suggested that dopaminergic and noradrenergic inputs are required for hippocampus-dependent LTM ^68,69^, which are likely activated by co-release of the two neuromodulators from LC axons ^70,71^.

We propose that astrocytic Gq signaling mediates the effect of novel experience to promote the persistence of long-term memory (**Sup. 8**). LC noradrenergic neurons fire phasically in response to novel and salient events ^72,73^ and high amounts of released NA in turn trigger astrocytic Gq signaling via α1ARs ^1,2^. Among possible PRPs for LTM, including Homer1a, Arc, BDNF and PKMζ (protein kinase Mζ), a recent study indicated that PKMζ in prefrontal cortex serves as a PRP for object recognition memory ^74^. Although astrocytic Gq signaling has yet been shown to induce PRPs in neurons, elevated adenosine A1R signaling in neurons is known to increase ERK-phosphorylation and Hormer1a expression *in vivo* ^75^. ATP release from astrocytes and the subsequent conversion to adenosine conceivably induce PRPs via adenosine A1R (➀ in **Sup. 8**).

Synaptic depression has been suggested as a key mechanism for novelty detection and memory ^76–78^. NA application to acute brain slices induced long-term depression (LTD) in an α1AR-dependent manner ^79,80^, which is presumably mediated by astrocytic ATP release ^81^. Moreover, phasic stimulations of LC *in vivo* induced βAR-dependent LTD in the hippocampus ^82^. Notably, serum response factor (SRF)-deficient mice are impaired of hippocampal LTD and do not habituate to novel environments ^83^. Our results of synaptic depression and reduced locomotion by astrocytic Gq activation (**Fig. 3 and 4**) complement these phenotypes and further postulate that astrocytic Gq activation may be involved in SRF signaling. Indeed, the α1AR has been shown to activate SRF in non-neuronal cells ^84^. Of particular note, a very recent study reported LTD deficiency in IP3R2-KO mice ^19^. Together, these findings lend support to the notion that novelty triggers astrocytic Gq signal and induces synaptic depression. Such a mechanism may underlie novelty-acquisition behaviors such as habituation to a novel environment. How LTD facilitates STM-to-LTM conversion remains unknown. It is tempting to speculate that LTP of relevant synapses for memory formation carries relatively high information under the LTD-favored condition, due to an improved signal-to-noise ratio.

## Materials and Methods

All experimental protocols were approved by the RIKEN Institutional Animal Care and Use Committee.

### Generation of transgenic mice

The PCR fragment of *Optoα1AR-EYFP* was ampified from the pcDNA3.1/Optoα1AR-EYFP plasmid ^27^(gift from Dr. Karl Deisseroth) and was subcloned into the FseI site of the Ai9 plasmid (Addgene). Then, the resultant *Optoα1AR-EYFP-WPRE-bGH* polyA fragment was amplified by PCR and was subcloned between the XhoI and EcoRV sites of the pCR-FRT-*Amp*-FRT plasmid ^85^(gift from Dr. Kunio Yaguchi). A bacterial artificial chromosome (BAC) clone, RPCI-23-361H22 ^28^(BAC PAC Resources), containing the *GLT1* gene was modified by the Red/ET recombination system (Gene Bridges) to insert *Optoα1AR-YFP*-WPRE-polyA-FRT-*Amp*-FRT to the immediate downstream of the initiation codon. After colony selection by ampicillin, the *Amp* cassette was removed from the recombinant BAC clones by introducing the Flp recombinase expression plasmid 706-FLP (Gene Bridges). The final BAC construct was amplified, purified with the Large-Construction kit (Qiagen) and linearized by AscI digestion. Correct modification of the BAC was verified by pulsed-field gel analysis of restriction digestions and direct sequencing of the insert. The linearized BAC DNA was purified, adjusted to be ~1 ng/μl in a microinjection buffer (10 mM Tris-Cl, 0.1mM EDTA, 100mM NaCl, pH 7.4), and individually injected into the pronuclei of 445 C57BL/6J-fertilized embryos. As a result, 81 founders were born, of which 11 founders were positive for the transgene. Positive founder mice were identified by 301 bp DNA amplified by PCR using the following primer pair: 5’ CGAGGCGCTAAAGGGCTTACC 3’ and 5’ CCCCAGCATAATCAGAAGGA 3’. Positive founder mice were crossed with C57BL/6J mice and maintained on this genetic background. The established 11 line gave astrocytic expression of Optoα1AR-EYFP with different expression strengths and variable positive cell proportion. Among them, the two lines, #941 and #877, were selected based on the selective expression of Optoα1AR-EYFP in astrocytes and used in the current study as strong TG and patchy TG mice, respectively. These heterozygous mice were crossed with C57BL/6J mice to obtain heterozygous mice for experiments and were noted as TG mice.

### Immunohistochemistry

Mice were transcardially perfused with 4% paraformaldehyde in 0.1 M phosphate buffer. Coronal sections of 60 μm thickness across AP ~-1.5 mm were obtained. These sections were incubated with anti-S100β (1:200, ab52642, Abcam), anti-NeuN (1:2000, MAB377, Chemicon-Millipore), anti-GFAP (1:2000, Z0334, DAKO) or anti-Iba1 (1:400, 019-19741, Wako) antibodies. Subsequently, sections were incubated with Alexa Fluor-594 conjugated secondary antibodies (1:1000, Molecular Probes-Invitrogen). Fluorescence images were obtained with a 10× (0.4 NA) or 60× (1.3 NA) objective lens using an Olympus FV-3000 confocal microscope.

### *In vivo* two-photon Ca^2+^ imaging of astrocytes

Adult Optoα1AR patchy TG mice (>2 months old) were anesthetized with 1.5 g/kg urethane. A metal frame was attached to the skull using a dental acrylic (Fuji LUTE BC, GC and Super Bond, Sun Medical), and a craniotomy (diameter 2.0-3.0 mm) for imaging was made above the somatosensory cortex (AP -1.5 to -2.5 mm and ML 1.5 to 2.5 mm). The dura mater was carefully removed, and the exposed cortex was loaded with Rhod-2 AM (0.4 mM, Molecular Probes-Invitrogen) for 1 hour. After washing with HEPES-ACSF (125 mM NaCl, 3.5 mM KCl, 10 mM glucose, 10 mM HEPES, 2 mM CaCl2 and 2 mM MgSO4, pH 7.3) several times, the craniotomy was covered with agarose (2% w/v in HEPES-ACSF) and sealed by a thin glass coverslip. A Bergamo based two-photon microscope (Thorlabs) attached to a Chameleon Ultra 2 laser (Coherent) with 25× objective lens (1.05 NA, Olympus) was used. The microscope is equipped with a reverse dichroic mirror (405/473-488/561/705-1600nm notched dichroic mirror, Thorlabs) and the emission light was separated by a dichroic mirror (FF562-Di03, Semrock), with band-pass filters FF03-525/50 and FF01-607/70 (both from Semrock) for the green and red channels, respectively. Rhod-2 fluorescence from 50–100 μm below the pial surface was acquired using ThorImageLS software at a frame rate of 5 Hz. Rhod-2 or EYFP was imaged with 820 nm or 940 nm laser, respectively.

Photo-activation for Optoα1AR was carried out by 470 nm LED (M470L3, Thorlabs) after systemic 9-cis-Retinal supply. Strong (1 mW) or weak (0.1 mW) LED was targeted to an imaged region (φ = ~0.8 mm) through the objective lens. To protect photomultiplier tubes (PMTs) from LED illumination, the optical path to the PMTs was blocked by a built-in mechanized shutter. 9-cis-Retinal (R5754, Sigma) was dissolved in Dimethyl sulfoxide (DMSO) to make 200 mM solution and stored frozen in 5 μL aliquots. On the day of experiment, an aliquot (i.e. containing 0.28 mg 9-cis-Retinal) was diluted in 100 μL HEPES-ACSF or saline at 35 °C and administered by intraperitoneal (i.p.) injection. This amount of 9-cis-Retinal is comparable to that in the previous study, where i.p. injection of 0.375 mg of 9-cis-Retinal restored electroretinogram responses in endogenous 11-cis-retinal deficient mice ^86^. LED-induced Rhod-2 responses were reliably observed ~30 min after 9-cis-Retinal injection and thereafter for ~1.5 hr. LED illuminations were repeated at ~9 min interval. Tail pinch was manually applied via blunt tongs for ~1 s.

For long-term Ca^2+^ imaging, RCaMP1.07 was selectively expressed in astrocytes under the control of a GFAP promoter. Optoα1AR strong TG mice (>2 months old) were anesthetized with ketamine and xylazine (56 and 8 mg/kg, respectively) and a metal frame was attached to the skull. A small craniotomy was made above the somatosensory cortex and a glass micropipette containing AAV9-hGFAP-RCaMP1.07 (3.0–4.0 × 10^12^ vg/ml, PBS) was inserted to a depth of 300 μm below the pial surface. Microinjection of 300 nl was made over 3 min using a Femtojet injector (~5 psi, Eppendorf), and the exposed cortical surface was covered by a sterilized round cover glass (3 or 4 mm in diameter) to be used as a cranial window for later imaging.

>2 weeks later, AAV-microinjected mice were anesthetized with 1.5 g/kg urethane, and RCaMP imaging was performed using the Bergamo two-photon microscope with 1040 - 60 nm laser. In these experiments, the same region was repeatedly imaged before and after LED illuminations (1 mW) with different durations (1 s, 3 s or 3 min) at 9 min interval. For 3 min illumination, 1.5 s LED-on and 0.5 s LED-off, were repeated. RCaMP signals were first imaged without 9-cis-Retinal addition. Thereafter, 9-cis-Retinal was supplemented by i.p. injection, and imaging was resumed after 40 min. For testing the repeatability of 1-s Optoα1AR activation, LED illuminations were repeated at 3-min intervals in addition to 9-min interval.

### *In vivo* two-photon imaging of blood vessels

Metal frames were attached to the skull of anesthetized adult Optoα1AR strong TG mice (>2 months old, 1.0 g/kg urethane and 50 mg/kg α-chloralose). A small craniotomy was made above the somatosensory cortex (AP -1.0 to 1.0 mm and ML 1.5 to 3.5 mm). The serum was labeled by intravenous injection of FITC-dextran (2 MDa, Sigma). Needles (30 G) were inserted in the contralateral forelimb to apply sensory stimulation (1 mA, 100 ms, 3 Hz, 6 times, interval 30 s, 10 times). Vasculature imaging was performed after 9-cis-Retinal i.p. injection using the Bergamo microscope (820 nm). Once sensory-driven vasodilation was observed, the same area was imaged with Optoα1AR activation by LED illumination (1 mW, duration 1s). In some mice (3 out of 5 mice), astrocytic Rhod-2 loading in the somatosensory cortex was also done. Simultaneous imaging of astrocytic endfoot Ca^2+^ and the vasculature was performed using the Bergamo microscope with 820 nm laser. For analysis of FITC area, penetrating arterioles that have circular cross-sections were sampled.

### *In vivo* two-photon Ca^2+^ imaging of neurons

AAV1.Syn.NES-jRGECO1a.WPRE.SV40 (Penn Vector Core, 3.4 – 5.7 x 10^12^ vg/ml) was injected at a depth of 300 μm of the somatosensory cortex of adult WT and Optoα1AR strong TG mice, as above. Following >2 weeks recovery, mice were imaged using the Bergamo microscope with 1040–60 nm laser at a frame rate of 5 Hz in anesthetized (1.5 g/kg urethane) or awake condition. In the latter case, mice were trained to be restrained under the microscope using a mechanical fixture that rigidly fixes the head frame once a day for 5–7 days. During this training, mice were water-deprived, and had access to water once the head frame is fixed to the apparatus, thereby making an association between the head fixture and satiation of thirst. In both anesthetized and awake conditions, 9-cis-Retinal i.p. injection was made ~40 min before imaging. Optoα1AR was activated by LED (1 mW, 1 s), which was repeated 4 to 8 times at 9-min interval. In some TG mice, adenosine A1R antagonist, CPT (C102, Sigma) was administered (20 mg/kg, i.p., dissolved at 40 mM in DMSO) 60 min before optogenetic stimulation.

### *In vivo* local field potential recording

Adult WT and Optoα1AR strong TG mice (>2 months old) were anesthetized with isoflurane (1.5%). A metal frame was attached to the skull and mice were rigidly fixed in a headplate holding device. A craniotomy was made above the somatosensory cortex. A glass micropipette (2 μm tip diameter, 1B150F-4, World Precision Instruments) filled with HEPES-ACSF was placed to an electrode holder with a headstage preamplifier. The headstage is then mounted to a remote-controlled micromanipulator (EMM-3NV, Narishige). Under a stereo microscope, the glass micropipette was inserted to the primary somatosensory cortex, trunk region (AP ~-1.5 mm, ML ~1.7 mm and DV ~0.25 mm) ^87^. An optical fiber (200 μm diameter, CFML22L10, Thorlabs) connected to 470 nm LED devices (LEDFRJ-B_FC and LEDRV_1CH_1000, Doric) was placed over the pial surface above the recording site. Two vitrodes (L150, Nihon Kohden) were put on the both sides of the dorsal trunk to apply sensory stimulation (1.5 mA, duration 1 ms, interval 10 s). After 9-cis-Retinal i.p. injection, isoflurane dosage was decreased to ~0.8 %. Thereafter, the room light was turned off.

After sensory evoked response was stabilized (typically ~1 hr), evoked field potential recording began (Multiclamp 700B, Axon instruments; 1000 x, 0.1 Hz to 3 kHz, digitized at 10 kHz using a LabVIEW-based data acquisition system, National Instruments). After stable evoked field potential response was obtained, brief LED illuminations were delivered above the recording site (1mW, duration 1 s, interval 5 min, 6 times).

### Behavioral experiments

Adult Optoα1AR TG mice and littermate WT mice (>2 months old) were anesthetized with ketamine and xylazine (56 and 8 mg/kg, respectively), and wireless LED device containing two LEDs (φ3.1, ~8.5mW x2) ^38^ was secured to skull above the anterior cortex (AP ~1.5 mm, ML ~1.5 mm) with dental cement. After >2-week recovery, mice were habituated to attachment of the LED-receiver and -battery with 9-cis-Retinal or vehicle i.p. injection for ~1-week (duration ~1 hr, interval 2-3 day, 4 times). This LED device attachment and systemic injection of retinal or vehicle were always done 40 to 60 min before the following behavioral tests. LED illumination began ~1 min after mice were put in the behavioral chambers.

#### Novel open field test

Individual mice were placed in the center of a novel open field (40 × 40 cm) and allowed to freely explore the arena for 45 min. The anterior cortex was transcranially illuminated by the LED device (duration 3 sec, interval 3 min, 15 times). In some TG mice, adenosine A1R antagonist, DPCPX (C101, Sigma) dissolved at 4 mg/mL in DMSO was i.p.-injected at the dosage of 1 mg/kg, 40 to 90 min before LED illuminations.

#### Y-maze test

Individual mice were placed in the center of a Y-maze (YM-40M, BrainScience idea) and allowed to freely explore the maze for 15 min with the LED illuminations (duration 3 sec, interval 3 min, 5 times) delivered.

#### Novel object recognition test

This test consisted of three different phases: habituation, training and test. Individual mice were first habituated to an open field with a plywood floor (39.5 × 39.5 cm), with 10 min of exploration 3 times in 3 consecutive days. The plywood floor was changed for each mouse, and the plywood floor was consistently used for each mouse in later training and test. On the training day, individual mice were placed in the habituated open field for 10 min, where two identical objects were fixed 20 cm apart on the plywood with double-sided adhesive tape (SRE-19, 3M). Three different objects of similar size were used in a counter-balanced manner, as the role (familiar vs novel object) as well as the position (left or right) were randomly permuted. Transient LED illuminations (duration 3 s, interval 3 min, 4 times) or longer LED illuminations (duration 30 s, interval 3 min, 4 times) were delivered during the training period. Immediately after training, mice were returned to the home cage, and LED-receiver and -battery were detached from mice 10 min later. In post-activation experiments, LED illuminations (duration 3 s, interval 3 min, 4 times) were delivered during this 10 min in the home cage immediately after training. On the test day, (1 day or 14 days after training), mice were exposed to one of the pre-familiarized objects and a novel object for 10 min without LED illumination. In some TG mice, DPCPX was i.p.-injected at the dosage of 1 mg/kg, ~60 min before LED illuminations.

#### Conditioned place preference test

The conditioned place preference apparatus consisted of three chambers. The left and right chambers have the same size (25 × 25 cm) but were distinguished by their walls with vertical-stripes or horizontal-stripes, respectively. The center chamber (25 × 15 cm) has neutral walls without stripe. This test consisted of three different phases: pre-test, conditioning and post-test. On day 1 (for pre-test), individual mice were placed in the center chamber and allowed to freely enter and explore the three chambers for 20 min without LED illumination to measure default place preference. On day 2 to 5 (for conditioning), mice pre-treated with 9-cis-Retinal were confined for 30 min to either the left or right chamber on alternate days, and LED illuminations (duration 3 s, interval 3 min, 10 times) were delivered in either the left or right chamber. Days with LED illuminations (day 2 and day 4 or day 3 and day 5) were counterbalanced. On day 6 (for post-test), mice were again placed in the center chamber and allowed to freely enter and explore the three chambers for 20 min without LED illumination to measure the post-conditioning place preference. Post-test and pre-test were conducted identically.

## Data analysis

### Immunohistochemistry

Quantifications (**Sup. 1**) were performed from layer 1 to 6 in both sides of 500 μm-wide somatosensory cortex (ML ~1.5-2.0 mm). Among the cellular marker-positive cells, EYFP-positive or EYFP-negative cells were manually counted from 60 x images at single plane. Intensity of GFAP, IbaI or EYFP of the above somatosensory cortical area was measured by ImageJ software from stack images acquired with a 10× objective over 60 μm-thick brain slices. Measured intensity of GFAP or IbaI of a mouse was normalized by mean intensity of WT mice and presented as relative intensity in **Sup. 1c** or **1d**. Measured intensity of EYFP of a TG mouse was first subtracted by mean intensity of WT mice which corresponds to background EYFP intensity. The subtracted value was then divided by the mean intensity of patchy TG mice and described as relative percent to patchy TG mice in the text.

### Two-photon imaging

Analysis was performed by ImageJ and MATLAB software. Image shift in xy axis was adjusted by the TurboReg ImageJ plug-in program for all images.

Rhod-2 and RCaMP signals in astrocytes (**Fig. 1 and Sup. 2**) were extracted from cell bodies manually on ImageJ, and these intensity data in region of interests (ROIs) were exported to MATLAB software for further analysis of F/F_0_, where F is fluorescence intensity within a given ROI at each time point and F_0_ is the mean fluorescence intensity within a given ROI during 0-1 min before LED illumination. A responsive cell was defined as a cell exhibiting >120 % F/F_0_ for >10 s within 30 s after LED illumination. Peak F/F_0_, onset time, peak time and offset time were analyzed for responsive cells. Onset time is the time firstly reaching 120 % F/F_0_. Offset time is a time firstly returning to 120 % F/F_0_ after peak. The values in each image were averaged across ROIs, and these normalized values were presented in **Fig. 1 and Sup. 2**.

FITC signals (**Fig. 2**) were binarized and circular cross-sections of penetrating arterioles were extracted using ImageJ. The binarized data were used to calculate the cross-section area using MATLAB. Astrocytic endfeet area was manually marked in the neighborhood of the penetrating arterioles and Rhod-2 signals were extracted accordingly using ImageJ. Rhod-2 signals were analyzed by MATLAB to calculate of F/F_0_. The values of FITC and Rhod-2 were averaged across imaging trials and normalized to the control period of 0-10 s before sensory stimulation or LED illumination. Mean ± SEM of the normalized trace is presented in **Fig. 2 c-f**. Mean of this normalized value from 0 to 5 s and this peak value from 0 to 20 s after sensory stimulation or LED illumination were presented in **Fig. 2 g** and **h**.

RGECO signals in neurons (**Fig. 3 and Sup. 3**) were extracted from cell bodies and neuropils manually on ImageJ, and these intensity data were analyzed by MATLAB to calculate F/F_0_ and standard deviation (std) of F/F_0_ in each ROI. These stds were averaged across ROIs and then further averaged across imaging trials. The mean std in each mouse was normalized by that in 0-1 min before LED illumination as 1, and this normalized value was presented as relative RGECO std in **Fig. 3** and **Sup. 3**.

### Local field potential recording

The slope of evoked LFP (**Fig. 3**) was calculated by MATLAB as described previously ^16,20^. First, the initial deflection of the LFP response was isolated. Next, the region for slope calculation was defined as the interval within 20 to 80% of the peak-to-peak amplitude of the negative deflection. The slope was computed by linear regression of the selected region. LFP slope within 1 min bin in each mouse was averaged with regard to 6 times LEDs, and this mean value was normalized by that in 0-1 min before LED illumination as 1, and this normalized value was presented as relative FP slope in **Fig. 3f**. Averaged LFP slope within 5 min bin in each mouse was normalized by that in 0-5 min before LED illumination as 1, and this normalized value was presented as relative FP slope in **Fig. 3g**.

### Behavioral experiments

The entire sessions in behavioral experiments were recorded by a video camera (C910, Logicool). Animal’s body position was determined by Any-maze behavior tracking software (Stoelting). Regarding novel open field test (**Fig. 4, Sup. 4 and Sup. 5**), time in the center zone (central 20 x 20 cm), total traveled distance and immobile time were calculated by Any-maze software. Speed of traveled distance in each mouse was averaged with regard to 15 times LEDs, and this mean value was normalized by that in 0-20 s before LED illumination as 1, and this normalized value was presented as relative speed in **Fig. 4f**. Regarding Y-maze test (**Fig. 5**), total number of arm entries and correct arm entry (the entry to the arm different from the current and immediate prior ones) were determined from the recorded video. Total traveled distance and immobile time were measured by Any-maze software. Regarding novel object recognition test (**Fig. 6 and Sup. 6**), contact time was defined as time in touching the objects with nose or forepaws, which was determined from the recorded video. Relative contact is a ratio of contact time to a replaced object (F2 or N in **Fig. 6**) to total contact time to both objects. Traveled distance was measured by Any-maze software. Regarding conditioned place preference (**Sup. 7**), times in the individual chambers were measured by Any-maze software. Relative time in LED chamber is a ratio of time in LED-illuminated chamber (**left chamber in Sup. 7**) to time in both left and right chamber.

## Statistics

All statistical analyses were performed by Prism software. Comparisons between two groups were analyzed by unpaired t-test. If variances were significantly different between them, Welch’s correction was applied (Welch’s t-test). For comparisons of data between before and after a manipulation, paired t-test was used. Comparisons of three or more groups were performed by one-way ANOVA with post-hoc Tukey’s test. If variances were significantly different between them, Kruskal-Wallis one-way ANOVA with post-hoc Dunn’ s test was applied. For two-factor comparisons, two-way ANOVA with post-hoc Bonferroni test was used. All values are expressed as mean ± SEM.

## Acknowledgements

This work was supported by the RIKEN Brain Science Institute, KAKENHI grants (23115522, 26117520), and HFSP (RGP0036/2014). We thank Prof. Maiken Nedergaard and members of the laboratory for comments on earlier versions of the manuscript. The authors declare no competing financial interests.

---

**Supplemental figure 1**

<u>“Strong” and “Patchy” TG mouse lines have astrocyte-selective Optoα1AR-EYFP expression without inflammation.</u>

(A and B) EYFP fluorescence images corresponding to the S100β and NeuN images in Fig. 1b and c are shown. Adjacent sections are also immunolabeled by microglial specific marker, IbaI. Among the cellular marker-positive cells, percent of EYFP-positive cells were quantified. Scale bar: 1 mm (left), 50 μm (the rest).

(C) GFAP immunoreactivity of WT, Strong TG, and Patchy TG mice. Cortical GFAP signals are weak and similar across the genotypes (somatosensory cortex: p>0.15, one-way ANOVA, 4 WT, 4 strong TG vs 4 patchy TG mice). Scale-bar: 1 mm.

(D) IbaI immunoreactivity of WT, Strong TG, and Patchy TG mice. IbaI signals are not increased in either of TG mouse lines (somatosensory cortex: p>0.16, one-way ANOVA, 4 WT, 4 strong TG vs 4 patchy TG mice). Scale-bar: 1 mm.

**Supplemental figure 2**

<u>Brief illumination of Optoα1AR fully and repeatably activates astrocytic Ca^2+^ elevation only after retinal addition.</u>

(A) Representative two-photon image of somatosensory cortex layer 2/3 of a urethane-anesthetized strong TG mouse with AAV-induced RCaMP expression in astrocytes (left). RCaMP signal was imaged with LED illuminations (1 mW) of varying durations (1 s, 3 s or 3 min) before and after retinal addition (i.p.). Scale bar: 50 μm.

(B and C) Ca^2+^ response of Optoα1AR-positive astrocytes upon 1-s or 3-min illumination (top and bottom, respectively), before and after retinal injection. 1 s or 3 min illumination failed to activate Ca^2+^ signaling before retinal injection. After retinal injection, 1-s illumination induced a transient Ca^2+^ elevation. The peak amplitude of Ca^2+^ elevation was similar for 3-min illumination, suggesting that the brief illumination induces the saturated amplitude of Ca^2+^ elevation. Small SEM (light-green) indicates a reproducible response.

(D) Before retinal injection, Ca^2+^ response was not detected in astrocytes upon 1 s, 3 s or 3 min illumination. Each symbol represents an individual imaging session (N=7, 4 and 5 sessions from 3 strong TG mice for 1 s, 3 s and 3 min illuminations, respectively).

(E) After retinal injection, responsive cells and peak F/F_0_ were observed upon illumination. These parameters were similar among varying illumination durations (p>0.51 and p> 0.14, one-way ANOVA and Kruskal-Wallis one-way ANOVA). Each symbol represents an individual imaging session (N=8, 7 and 6 from 3 strong TG mice for 1 s, 3 s and 3 min illuminations, respectively).

(F) Demonstration of repeatability of Optoα1AR stimulation. One-minute interval results in a notable attenuation of Ca^2+^ response. Individual and mean Ca^2+^ responses upon 9 times LED illuminations were indicated by gray and green traces, respectively (26 astrocytes).

(G) After retinal injection, imaging was performed at intervals of 9 min and 3 min as indicated in A. Proportion and response amplitude of responsive cells were similar for both inter-stimulus intervals (9 min: p>0.29 and p>0.07, paired t-test, 4 pairs; 3 min: p>0.12 and p>0.09, paired t-test, 4 pairs). As the values induced by the 1st and 2nd illuminations were not significantly different, these were mixed and presented in (E).

**Supplemental figure 3**

<u>Brief astrocytic Gq activation under urethane-anesthesia did not significantly suppress spontaneous neuronal activity.</u>

(A) Relative Ca^2+^ activities of neuronal somata and neuropil are calculated as the standard deviation (std) of RGECO F/F_0_ in urethane-anesthetized strong TG mice. Somata or neuropil Ca^2+^ activities were not affected by LED illumination under urethane-anesthesia (soma post-LED 1-min period: 95.3±2.7%; p>0.17, paired t-test vs pre-LED 1-min period; neuropil post-LED 1-min period: 92.2±3.4%; p>0.10, paired t-test vs pre-LED 1-min period, 4 mice).

(B) Baseline Ca^2+^ activities of neuronal somata and neuropil were visibly lower in the urethane-anesthetized condition than in awake conditions (soma awake pre-LED 1-min period: 204.8±11.9% relative to soma urethane pre-LED 1-min period; p<0.001, unpaired t-test; neuropil awake pre-LED 1-min period: 198.8±13.9% relative to neuropil urethane pre-LED 1-min period; p<0.001, unpaired t-test).

**Supplemental figure 4**

<u>Patchy TG mice show a trend of decreased open-field activity during LED activations.</u>

(A) Novel open field test of WT and patchy TG mice during transient LED illuminations (duration 3 sec, interval 3 min, 15 times) with retinal pre-treatment. Representative locomotion traces and time maps for 45 min show that the TG mouse traveled shorter distances, while spending a similar length of time in the center zone as the WT mouse. Color bar: 15 s.

(B) Time in center zone was not significantly different between genotypes and between 15 min periods (p>0.53 and p>0.42, two-way ANOVA, 5 WT mice vs 6 patchy TG mice).

(C) TG mice tended to travel a shorter distance (*p<0.05, unpaired t-test, 5 WT mice vs 6 patchy TG mice).

(D) TG mice tended to have increased immobile time (**p<0.01, *p<0.05, unpaired t-test, 5 WT mice vs 6 patchy TG mice).

**Supplemental figure 5**

<u>Decreased open-field locomotion activity in strong TG mice by LED activations depends on retinal addition.</u>

(A) Novel open field test of WT and strong TG mice during transient LED illuminations (duration 3 sec, interval 3 min, 15 times) with pre-treatment of vehicle instead of retinal. Representative locomotion traces and time maps for 45 min show that the TG mouse traveled similar distance and spent similar time in the center zone as the WT mouse. Color bar: 15 s.

(B) Time in center zone was not significantly different between genotypes and between 15 min periods (p>0.81 and p>0.57, two-way ANOVA, 5 WT mice vs 6 strong TG mice).

(C) TG mice traveled similar distance as WT mice (p>0.24, unpaired t-test, 5 WT mice vs 6 strong TG mice).

(D) TG mice had similar immobile time as WT mice (p>0.11, unpaired t-test, 5 WT mice vs 6 strong TG mice).

**Supplemental figure 6**

<u>Astrocytic Gq activation immediately after training enhances long-term memory, and exploratory behaviors during training is not related to memory effects.</u>

(A) Strong TG mice received LED illuminations (duration 3 s, interval 3 min, 4 times) in the home cage immediately after training. This post-activation protocol also induced the novel object preference 14 days after training (p<0.02, paired t-test, 11 strong TG mice).

(B) Distance traveled during training was not significantly different between genotypes, between LED durations or between drug treatments (p>0.54, unpaired t-test, 21 WT mice with 3 s LED vs 22 TG mice with 3 s LED; p>0.28, one-way ANOVA, 22 TG mice with 3 s LED, 11 TG mice without LED vs 9 TG mice with 30 s LED; p>0.06, one-way ANOVA, 22 TG mice with 3 s LED, 8 TG mice with 3 s LED and without retinal vs 18 TG mice with 3 s LED and with DPCPX).

(C) Total contact time with both objects during training was not significantly different between genotypes, between LED durations or between drug treatments (p>0.94, unpaired t-test, 21 WT mice with 3 s LED vs 22 TG mice with 3 s LED; p>0.12, one-way ANOVA, 22 TG mice with 3 s LED, 11 TG mice without LED vs 9 TG mice with 30 s LED; p>0.79, one-way ANOVA, 22 TG mice with 3 s LED, 8 TG mice with 3 s LED and without retinal vs 18 TG mice with 3 s LED and with DPCPX).

**Supplemental figure 7**

<u>Anterior cortical astrocytic Gq activation does not affect conditioned place preference.</u>

(A) On day 1, mice freely entered and explored in the three chambers for 20 min to measure default place preference (pre-test). The left and right chambers were distinguished by their walls with vertical-stripes or horizontal-stripes, respectively. On day 2 to 5, mice pre-treated with retinal were confined for 30 min to either the left or right chamber on alternate days, and LED illuminations (duration 3 s, interval 3 min, 10 times) were delivered in either the left or right chamber (conditioning). Days with LED illuminations (day 2 and day 4 or day 3 and day 5) were counterbalanced. On day 6, mice again freely entered and explored in the three chambers for 20 min to measure the post-conditioning place preference (post-test).

(B) Time in each chamber was not significantly different between pre- and post-test in WT mice (p>0.99, two-way ANOVA, 5 WT mice).

(C) Time in each chamber was not significantly different between pre- and post-test in TG mice (p>0.99, two-way ANOVA, 7 strong TG mice).

(D) Relative time in LED chamber was not changed between pre- and post-test in WT mice (p>0.96, paired t-test).

(D) Relative time in LED chamber was not changed between pre- and post-test in TG mice (p>0.55, paired t-test).

**Supplemental figure 8**

<u>Proposed model for astrocytic Gq signaling-mediated enhancement of long-term memory in novelty detection.</u>

Novel experience induces synaptic depression, behavioral habituation, and LTM enhancement (all outlined with red). Our findings indicate that optogenetic activation of astrocytic Gq signaling can strengthen these novelty-induced effects through A1R signaling. Events are usually memorized only for a short time, unless they contain sufficient novelty to activate LC. When novelty is increased by novel open-field exposure, the resultant LC phasic activity converts STM to LTM, which likely involves β adrenergic receptors and D1/D5 dopaminergic receptors. According to the “synaptic and behavioral tagging” hypothesis, event-related neuronal activity sets “learning tags” in the relevant cells and synapses, and novelty-induced LC activity synthesizes “plasticity-related proteins (PRPs)”, which are captured by learning tags to promote memory consolidation. Astrocytic Gq signaling induced by LC activity can contribute to the long-term memory enhancement presumably through A1R signaling-dependent PRPs synthesis (➀) and/or synaptic depression-related mechanisms (➁).

**Supplemental movie 1**

<u>Astrocytic Ca^2+^ imaging upon brief Optoα1AR activation by weak LED illumination, corresponding to the Fig. 1F.</u>

Rhod-2 F/F_0_ movie was overlaid on a static image of Optoα1AR-EYFP (green) and Rhod-2 (red). White circles and arrowheads at the beginning of the movie indicate EYFP-positive and negative astrocytes, respectively. Weak LED illumination (0.1 mW, 1 s, blue screen at time 0–1 s) increases Ca^2+^ in EYFP-positive astrocytes, but not in EYFP-negative astrocytes. Scale bar: 50 μm; Color bar: 300 % F/F_0_.

**Supplemental movie 2**

<u>Astrocytic Ca^2+^ imaging upon brief Optoα1AR activation by strong LED illumination, corresponding to the Fig. 1G.</u>

Rhod-2 F/F_0_ movie was overlaid on a static image of Optoα1AR-EYFP (green) and Rhod-2 (red). White circles and arrowheads at the beginning of the movie indicate EYFP-positive and negative astrocytes, respectively. Strong LED illumination (1 mW, 1 s, blue screen at time 0–1 s) increases Ca^2+^ in EYFP-positive astrocytes, which appears to propagate to EYFP-negative astrocytes. Scale bar: 50 μm; Color bar: 300 % F/F_0_.

**Supplemental movie 3**

<u>Neuronal Ca^2+^ imaging upon brief Optoα1AR activation, corresponding to Fig. 3A.</u>

(Upper left) Static image of Optoα1AR-EYFP (green) and jRGECO1a (red). White rectangle indicates the analyzed area in Fig. 3A. Scale bar: 100 μm.

(Upper right) jRGECO1a F/F_0_ movie. White rectangle at the beginning of the movie indicates the same area as Fig. 3A. LED illumination (1 mW, 1 s, blue screen at time 0–1 s) induces a rapid neuronal Ca^2+^ decrease in the entire field of view, on which LED has been illuminated through the objective lens.

(Bottom) jRGECO1a F/F_0_ traces of cell 1-5 indicated in Fig. 3A are synchronized with the movie. Scale bars: 50 % F/F_0_ and 15 s.

