## Supplementary figures and images for "Transient astrocytic Gq signaling underlies remote memory enhancement"

### Sup. 1

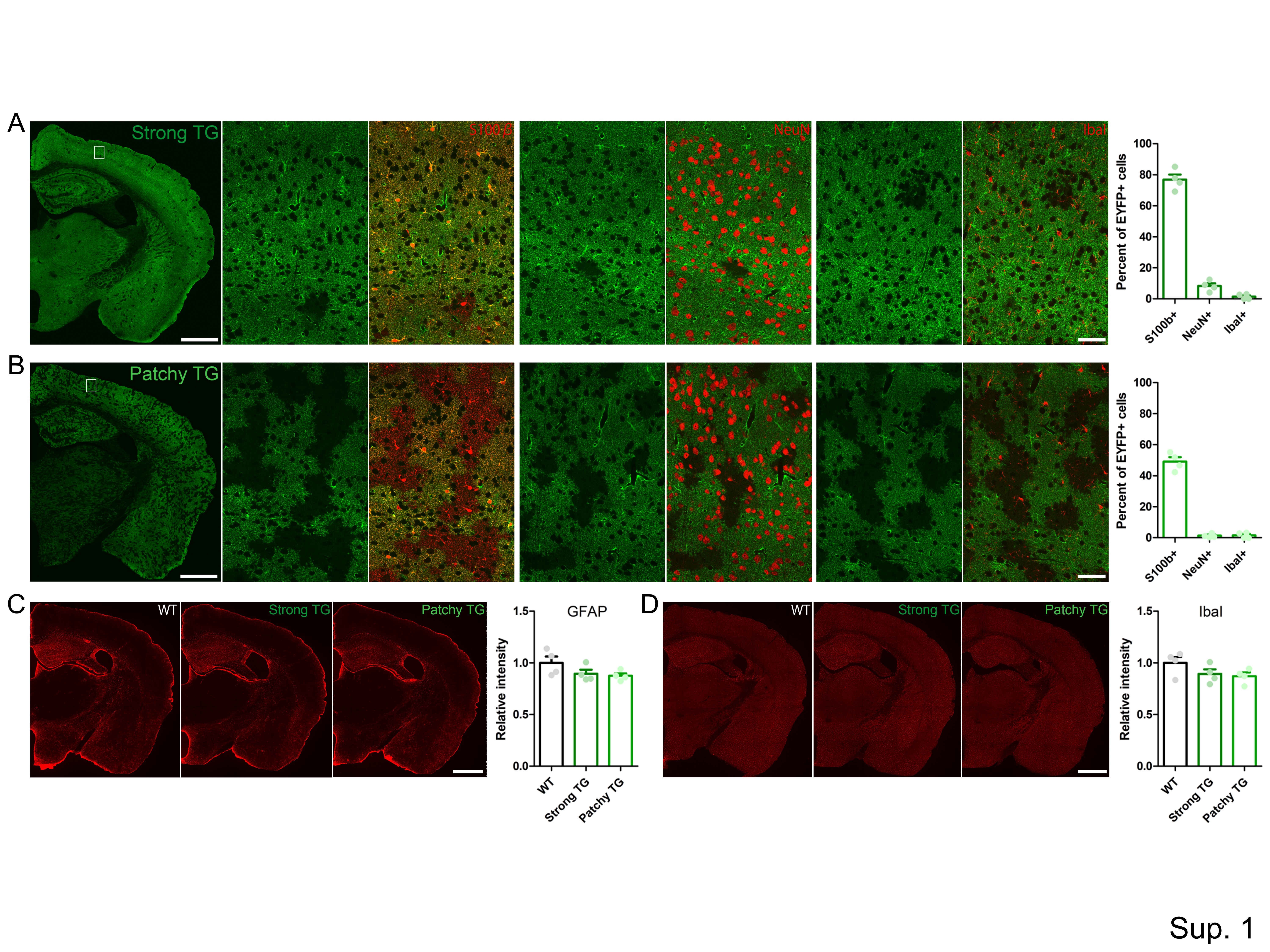

### Sup. 2

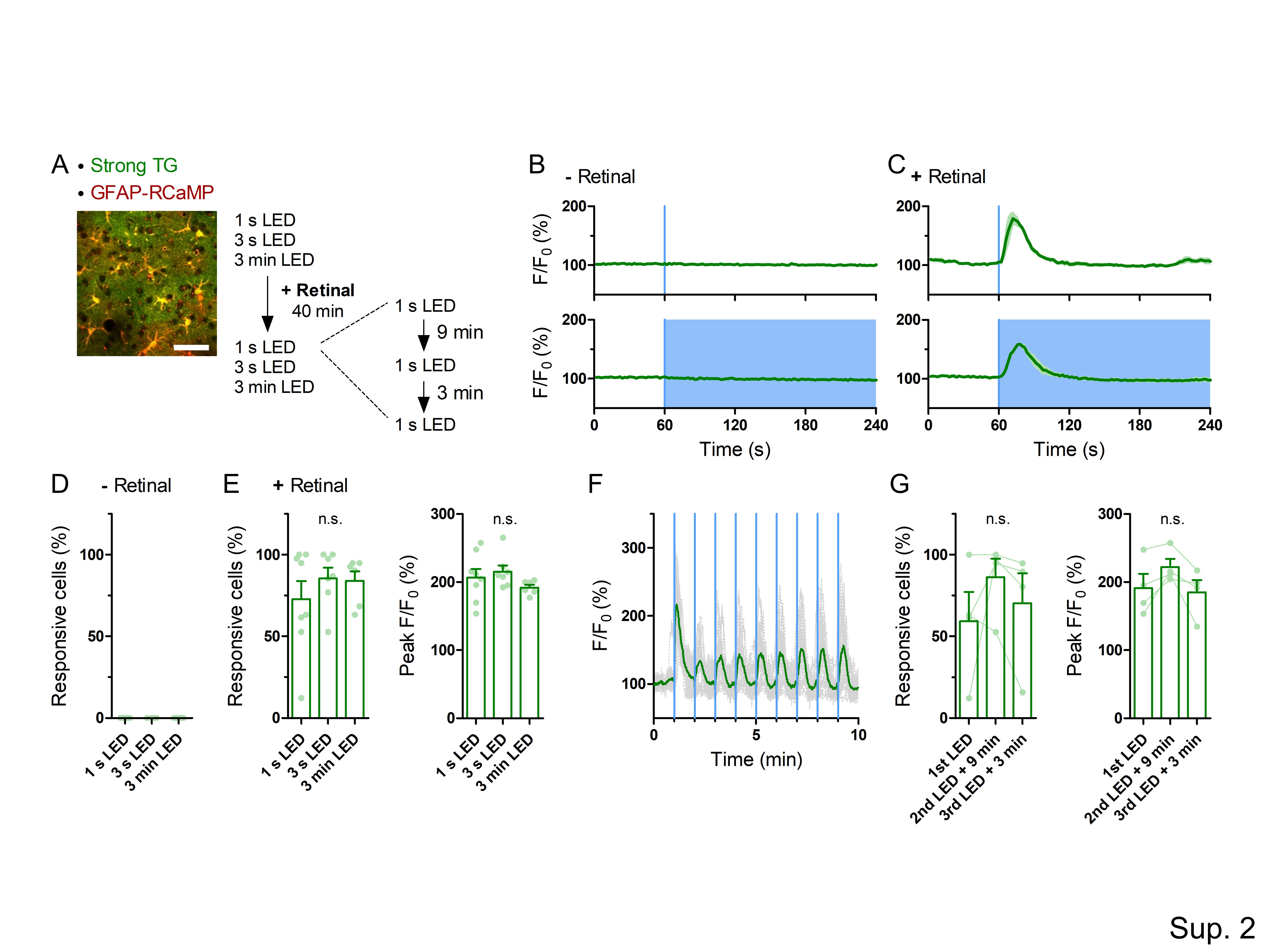

### Sup. 3

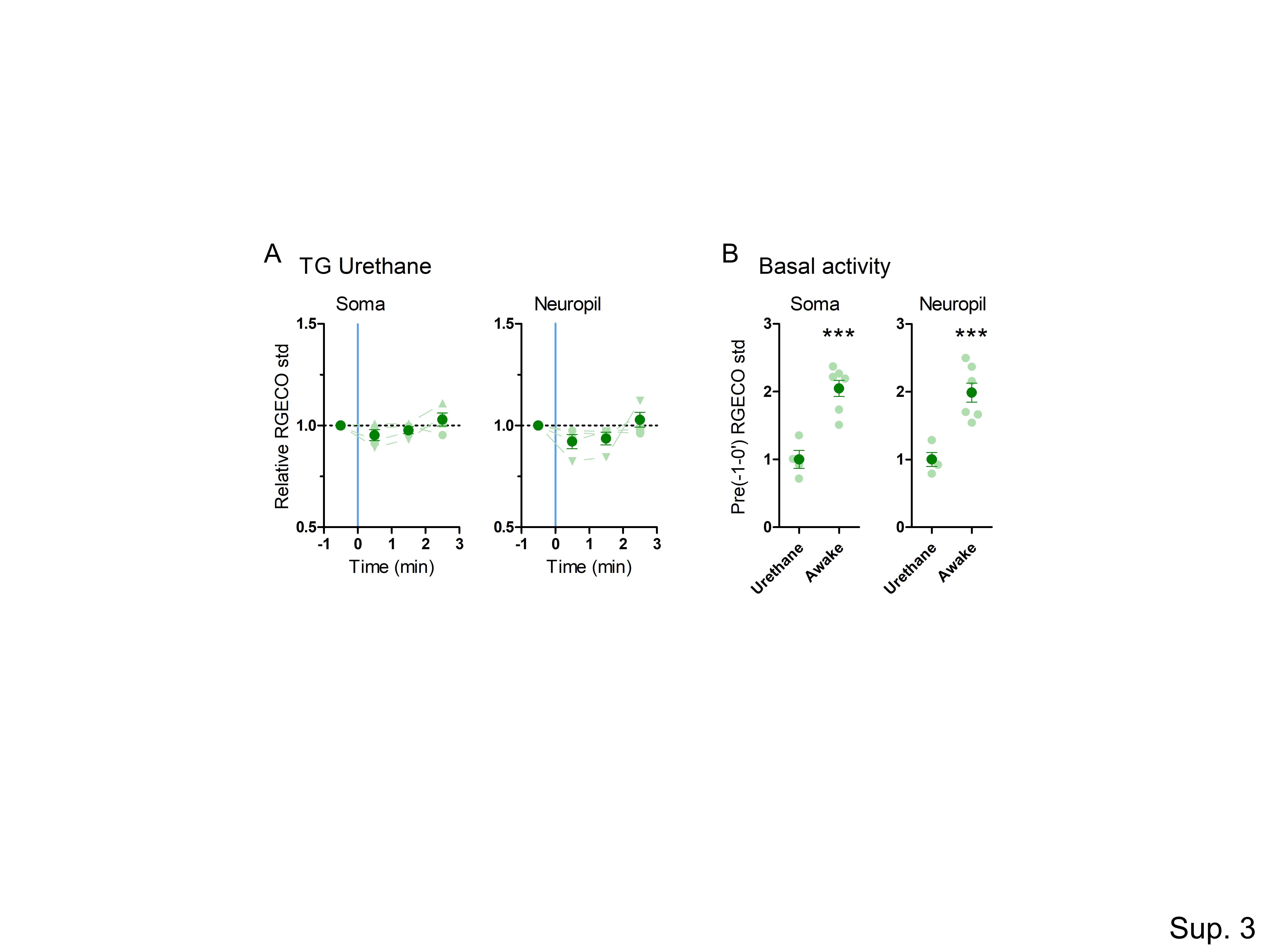

### Sup. 4

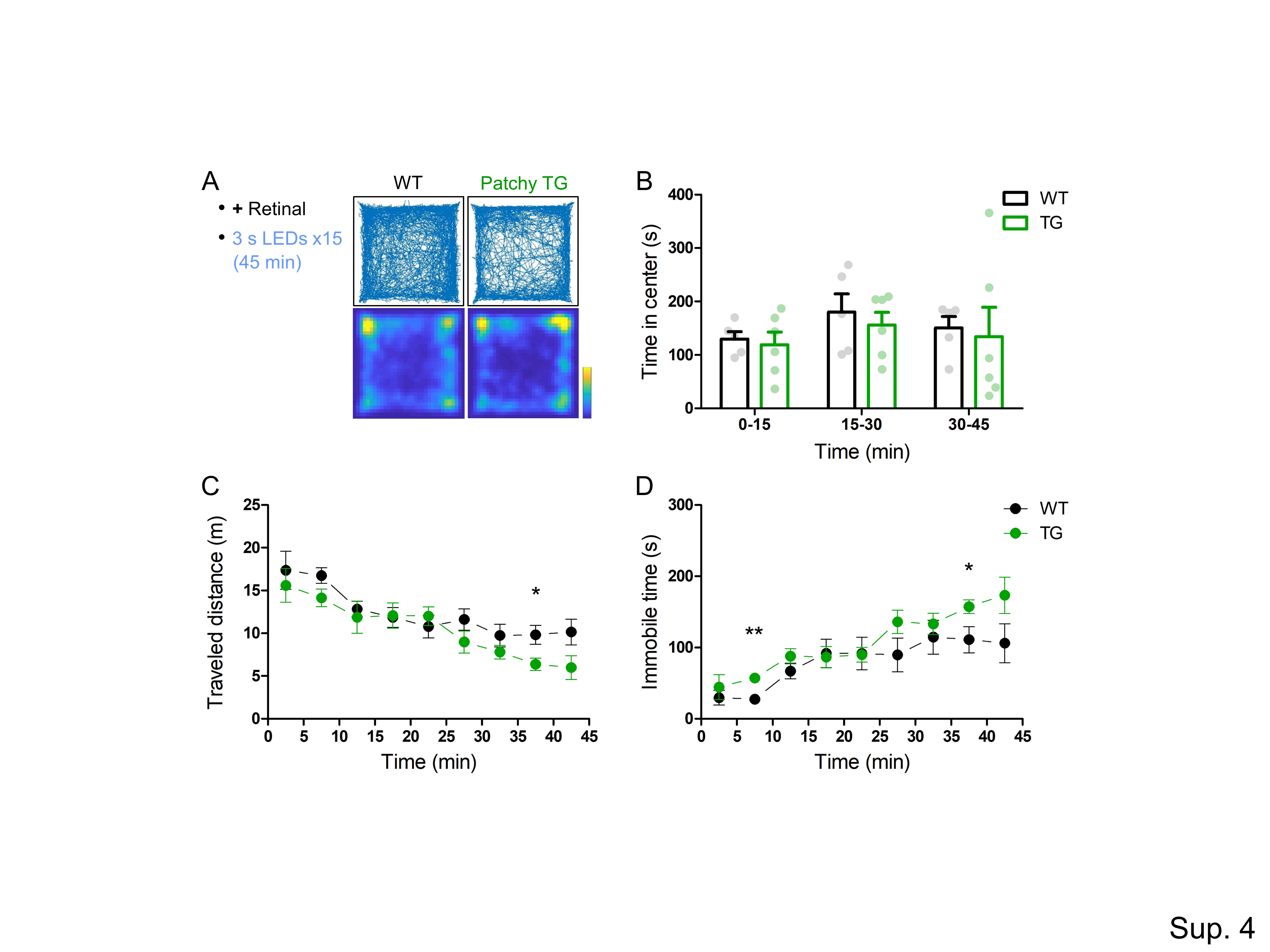

### Sup. 5

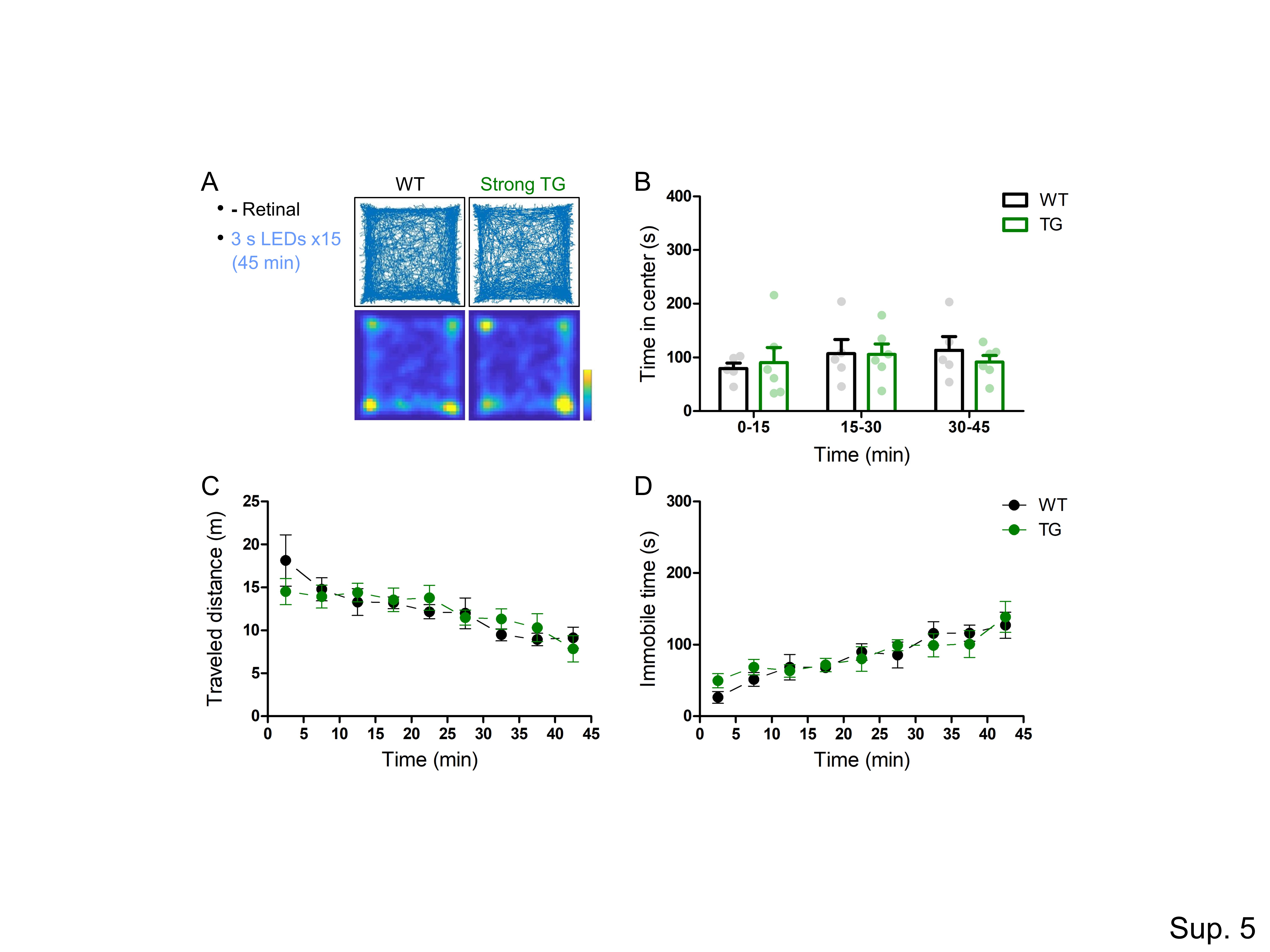

### Sup. 6

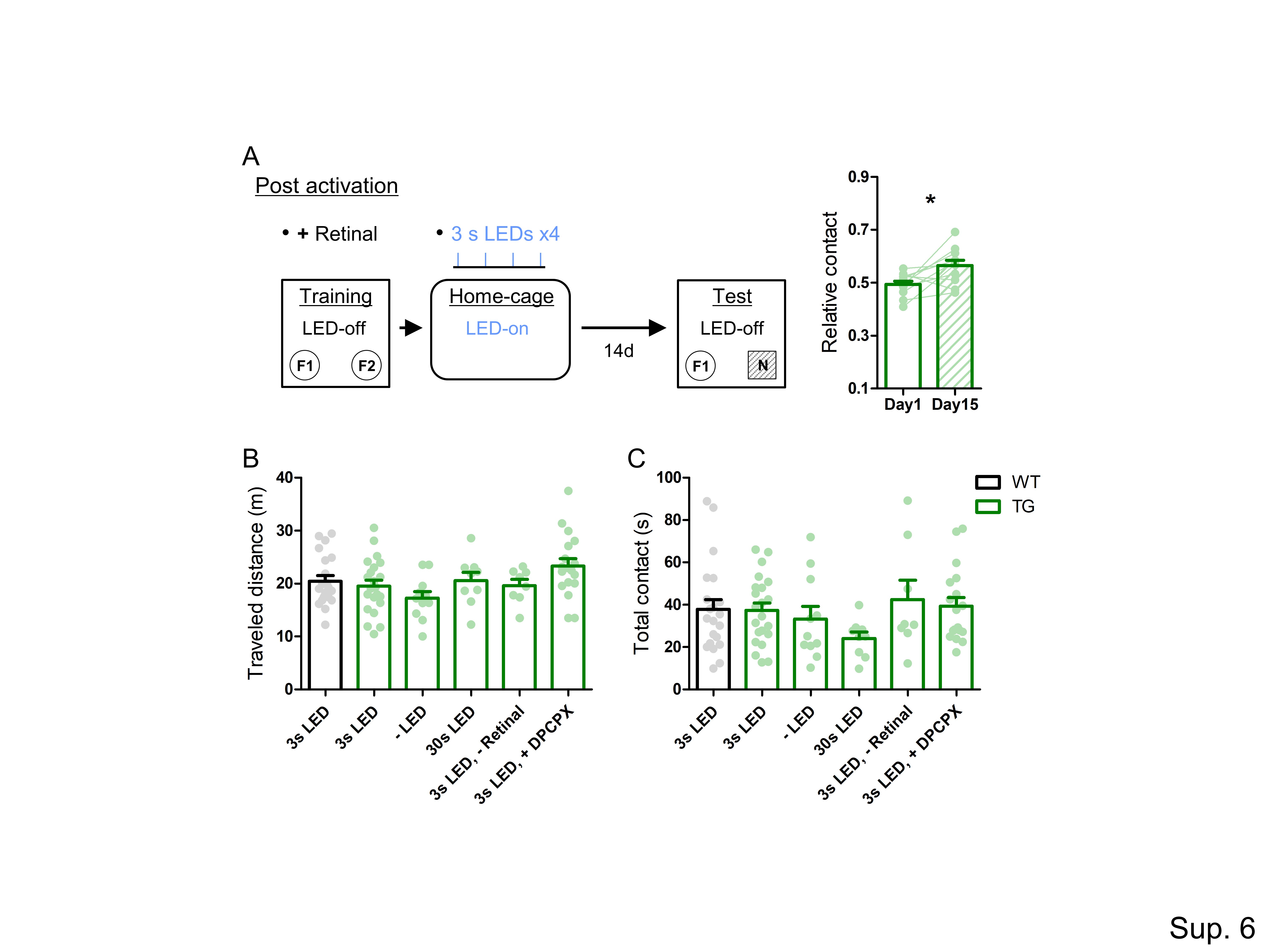

### Sup. 7

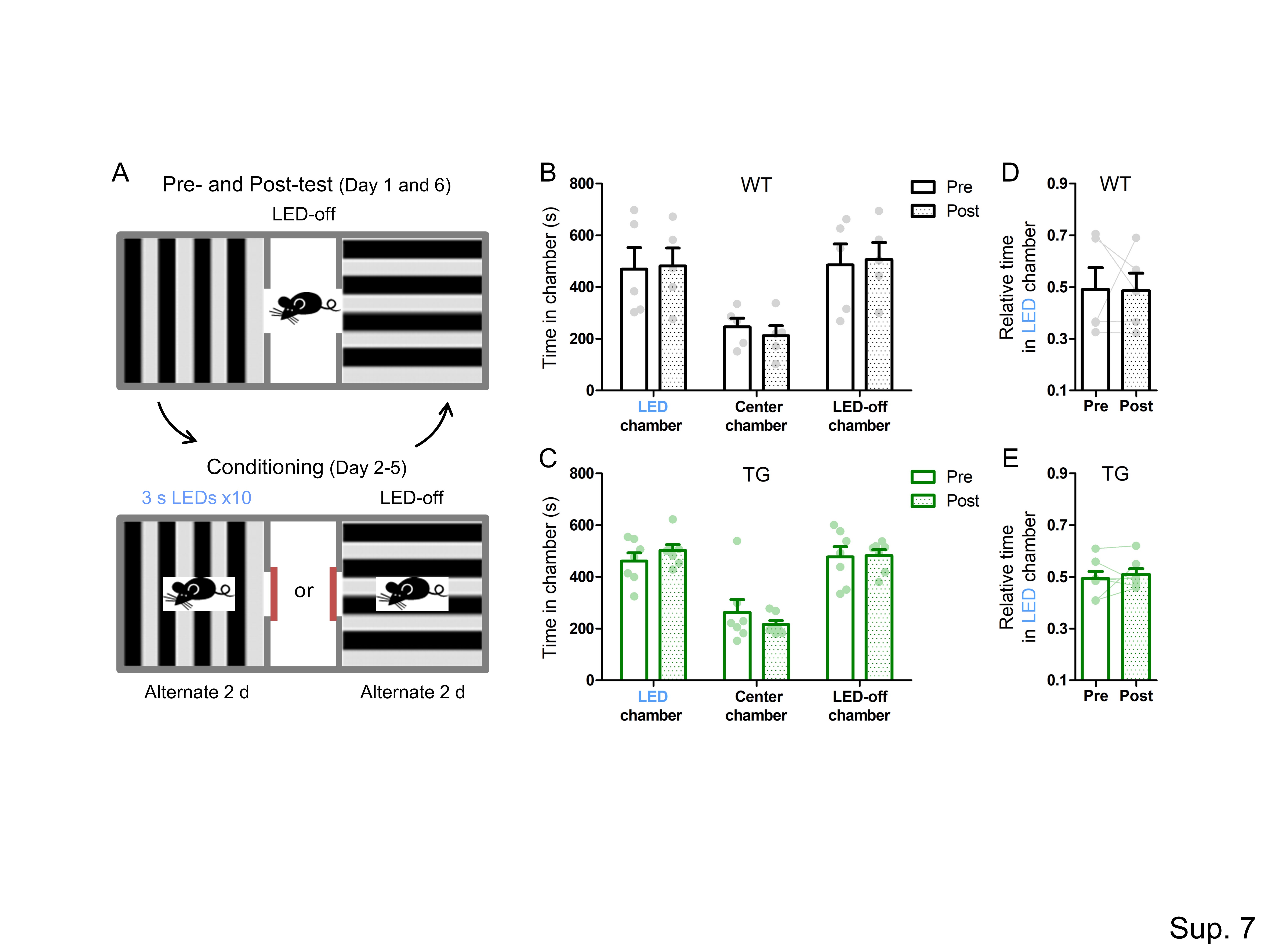

### Sup. 8

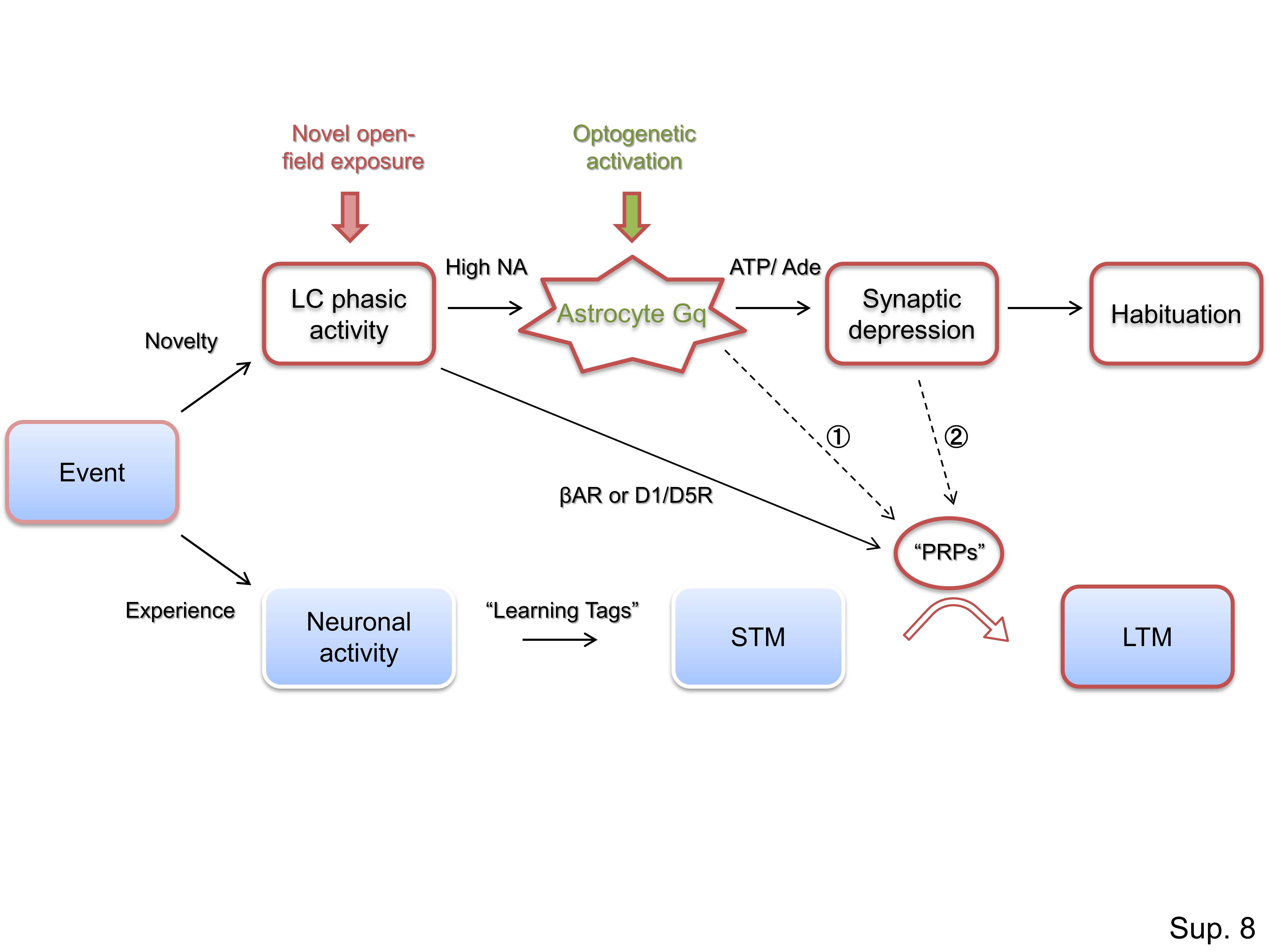
